# Genetic contributions of noncognitive skills to academic development

**DOI:** 10.1101/2023.04.03.535380

**Authors:** Margherita Malanchini, Andrea G. Allegrini, Michel G. Nivard, Pietro Biroli, Kaili Rimfeld, Rosa Cheesman, Sophie von Stumm, Perline A. Demange, Elsje van Bergen, Andrew D. Grotzinger, Laurel Raffington, Javier De la Fuente, Jean-Baptiste Pingault, K. Paige Harden, Elliot M. Tucker-Drob, Robert Plomin

## Abstract

Noncognitive skills such as motivation and self-regulation, are partly heritable and predict academic achievement beyond cognitive skills. However, how the relationship between noncognitive skills and academic achievement changes over development is unclear. The current study examined how cognitive and noncognitive skills contribute to academic achievement from ages 7 to 16 in a sample of over 10,000 children from England and Wales. Noncognitive skills were increasingly predictive of academic achievement across development. Twin and polygenic scores analyses found that the contribution of noncognitive genetics to academic achievement became stronger over the school years. Results from within-family analyses indicated that associations with noncognitive genetics could not simply be attributed to confounding by environmental differences between nuclear families and are consistent with a possible role for evocative/active gene-environment correlations. By studying genetic effects through a developmental lens, we provide novel insights into the role of noncognitive skills in academic development.

## Introduction

Children who are emotionally stable, motivated, and capable of regulating their attention and impulses do better in school, independent of their level of cognitive ability^1–7^. These important socioemotional characteristics have been broadly described as *noncognitive skills*^8^. “Noncognitive” is an imperfect term that primarily serves to differentiate these characteristics from what they are not – performance on standardized tests of cognitive ability. The panoply of noncognitive skills that predict better educational outcomes can be organized into three partly overlapping domains: motivational factors, self-regulatory strategies, and personality traits^9^.

Twin research has shown that genetic differences between people contribute to their differences in noncognitive skills. Most domains of noncognitive skills, including academic motivation^10,11^, self-regulation^12^ and personality,^13^ are moderately heritable (∼30-50%). In addition, twin studies have found evidence that noncognitive skills are genetically correlated with academic achievement^14,15^. That is, some of the same genetic differences that are associated with variation in academic achievement are also associated with noncognitive skills.

DNA-based methods have confirmed genetic links between noncognitive skills and academic performance. Genome-wide association studies (GWAS) of educational attainment (*i.e.*, years of formal education completed) have identified genetic variants that are correlated with completing formal education^16,17^. A polygenic score (PGS) constructed from these GWAS results predicts higher levels of self-control^18^, more adaptive personality traits (higher conscientiousness, agreeableness, and openness to experience), and greater academic motivation^19^. Additionally, previous GWAS work has identified associations between DNA variants and educational attainment that were independent of cognitive test performance, essentially performing a GWAS of noncognitive skills^20^. The genetics of noncognitive skills were found to be related to conscientiousness, openness to experience, delay of gratification, and health-risk behaviours^20^.

The current study uses both twin and DNA-based methods to expand our understanding of the role of noncognitive skills in academic development. We address four key questions (Figure 1). First, does the contribution of noncognitive skills to academic achievement change over development (from age 7 to age 16)? Second, do genetic predispositions to noncognitive skills vary in their contributions to academic achievement across development? Third, to what extent are these associations accounted for by between-family processes, such as environmental influences shared between individuals in a family? Fourth, do genetic contributions to academic achievement vary by socioeconomic status?

**Figure 1.**
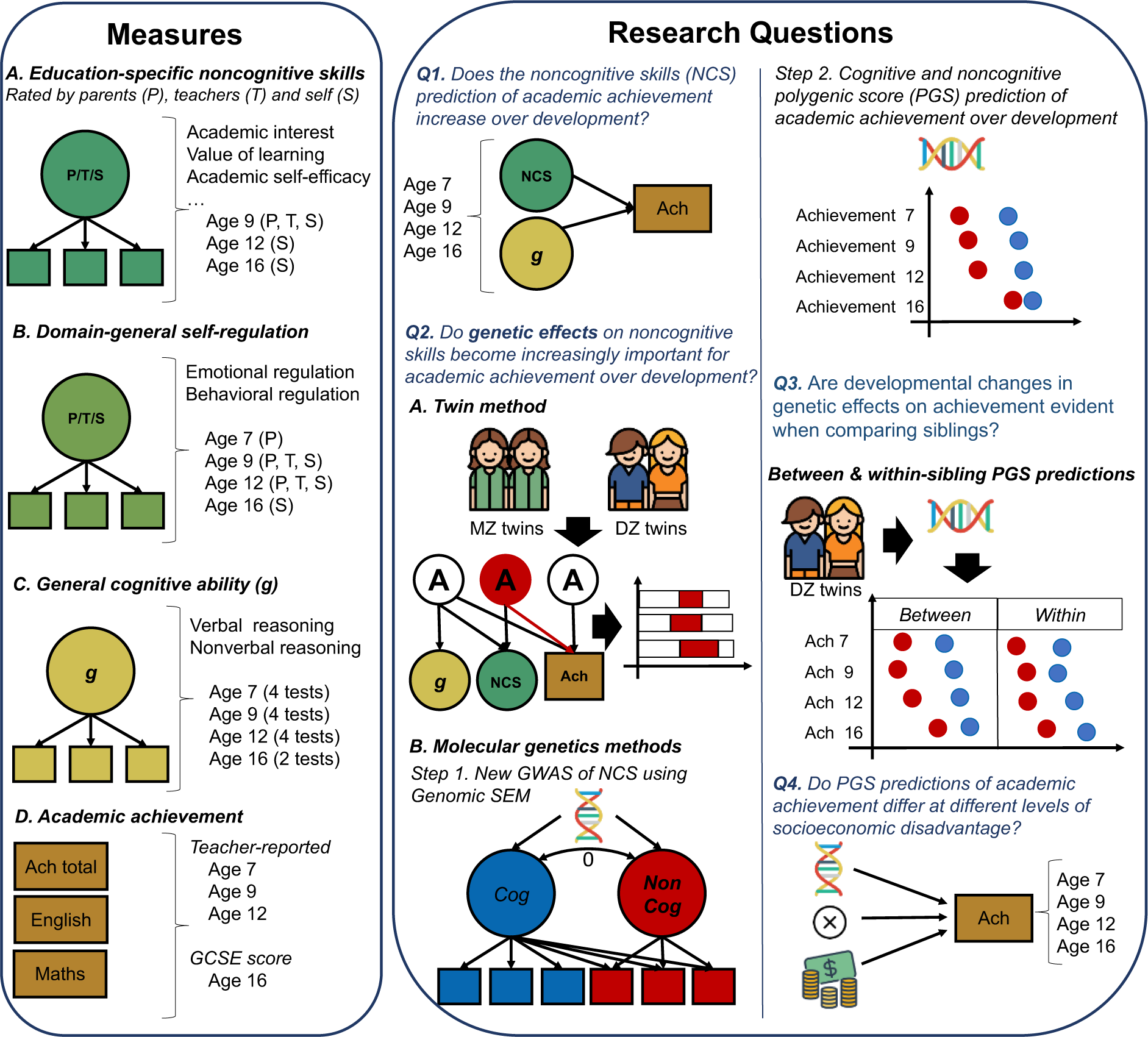
A visual summary of the measures, research questions and methods adopted in the present study. **Left panel:** We used factor analysis to capture individual differences in two broad dimensions of noncognitive skills: <u>education-specific noncognitive skills</u> (including measures such as academic interest, academic self-efficacy and value attributed to learning), and <u>domain-general self-regulation skills</u> (including measures of behavioural and emotional regulation not necessarily related to the school context). We also created latent measures of general cognitive ability from verbal and nonverbal cognitive tests at four ages. Academic achievement measures included teacher ratings of academic performance based on the national curriculum at ages 7, 9 and 12 and exam scores at age 16 (see Methods for a detailed description). **Centre and right panels:** A summary of the methodologies adopted to address each of the four core research questions in the study. We addressed the first research question (Q1) by conducting a series of multiple regressions to investigate changes in the developmental contribution of noncognitive skills to academic achievement beyond cognitive skills. We addressed the second research question (Q2) using multiple genetic methods. First (A), we conducted trivariate Cholesky decompositions using twin data. Second (B), we created a new GWAS of noncognitive skills by extending the GWAS-by-subtraction (Demange et al., 2021) approach with a set of GWAS for specific cognitive tasks and SES-relevant traits and examined developmental changes in the cognitive (Cog) and noncognitive (NonCog) polygenic score prediction of academic achievement from age 7 to 16. We addressed our third research question (Q3) by modelling Cog (blue) and NonCog (red) PGS effects within a sibling difference design, therefore separating within-family from between-family effects. We investigated our fourth research question (Q4) fitting multivariable models including the effects of the Cog/NonCog PGS, family socioeconomic status, and their two-way interaction.

First, we investigated the associations between noncognitive skills and academic achievement across development. Longitudinal studies that have examined the contribution of noncognitive skills to academic achievement remain scarce and have focused on a few specific measures over relatively short time frames^21^. Here, we analyze a comprehensive battery of developmental data from over 10,000 children born in England and Wales who were followed across compulsory education (Figure 1, left panel). Moreover, we simultaneously consider the role of cognitive skills. Past research has highlighted how skills that are broadly considered noncognitive, such as self-control, rely on cognitive competencies.^22^ Therefore, it is important to consider the role of developing cognitive skills when assessing the relationship between noncognitive skills and academic achievement over time.

Second, we investigated whether genetic dispositions towards noncognitive skills become increasingly important for academic achievement across development. Twin studies focusing on specific moments in childhood^23^ or adolescence^24^ have found that heritable variation in noncognitive skills, such as motivation and self-regulation, contribute to academic achievement beyond cognitive skills^25^. However, to our knowledge, no study to date has examined this relationship across development. We triangulate evidence across different methods, including twin and PGS analyses, to investigate the contribution of genetic factors associated with cognitive and noncognitive skills to academic achievement.

Third, with a sibling-difference design, we examined to what extent the developmental relationship between genetic propensity for noncognitive skills and academic achievement was accounted for by family-wide environmental processes. Sibling differences in genotypes are randomized by meiosis, such that siblings have an equal probability of inheriting any given parental allele. Therefore, within-sibling pair PGS associations are thought to be less confounded by environmental differences between nuclear families, including population stratification and indirect genetic effects ^26^. Indirect genetic effects refer to the effects of the non-transmitted parental genotypes on the offspring phenotype, potentially reflecting rearing environments, although they can also capture broader demographic phenomena, such as assortative mating^27^.

Conversely, differences between siblings in PGS associations are often referred to as “direct” genetic effects^28,29^ in that they are consistent with a causal effect of genetic variants within an individual on their phenotype. However, even direct genetic effects involve mediation through environmental processes. For example, children with a greater motivation towards academic achievement might actively select, modify, and create environmental experiences that foster further achievement, such as deciding to take advanced classes^29^. That is, genetic differences between children can result in differential exposure to learning environments, which, in turn, can affect their academic achievement, ^30^. These active/evocative gene-environment correlations (rGE) amplify the effects of genetic difference and are one theorized mechanism for increasing genetic effects over development^31,32^.

Fourth, we explored whether genetic contributions to academic achievement varied by socioeconomic status. Genetic and environmental processes might interact such that the effects of environmental experiences on a trait might be partly dependent on genetic effects and vice versa.^33,34^ Studies that examined this possibility have focused on the role of socioeconomic disadvantage across a broad range of contexts, including family socioeconomic status^35,36^ and the school environment^37,38^. We explore whether the cognitive and noncognitive PGS prediction of academic achievement differs at different levels of socioeconomic disadvantage across development.

Under a developmental lens, these analyses address four core research questions providing a detailed account of the processes through which cognitive and noncognitive skills are linked to individual differences in academic achievement. We triangulated evidence across multiple genetic methods.

Since each method is subject to different and unrelated assumptions and limitations, triangulating multiple methods provides a powerful tool to increase the reliability of our results.

## Results

### Noncognitive skills predict academic achievement beyond cognitive skills with increasingly strong associations across development

Parents, teachers, and twins rated different noncognitive skills at different ages. Based on extant literature and measures availability, we focused on two broad dimensions of noncognitive skills, which were modeled as latent factors (Figure 1): 1) education-specific noncognitive skills, including measures of academic interest, attitudes towards learning, and academic self-efficacy, and 2) domain-general self-regulation skills, including measures of behavioural and emotional regulation not necessarily related to the school context (Figure 1 and Methods). Here, we report analyses of these two dimensions. Analyses of individual measures are reported in the Supplementary Material (Supplementary Note 1, Supplementary Figure 1, and Supplementary Tables 1 and 2).

Latent factors of education-specific noncognitive skills and domain-general self-regulation skills (Supplementary Tables 3 and 4) were correlated positively with academic achievement at all developmental stages. Effect sizes differed by rater and developmental stage and tended to increase with age. For example, the association between self-rated education-specific noncognitive skills and academic achievement increased from *r* = 0.10 (95% CIs = 0.07; 0.14) at age 9, to *r* = 0.41 (95% Cis = 0.38; 0.44) at age 12, to *r* = 0.51 (95% CIs = 0.48; 0.55) at age 16 (see Supplementary Note1, Supplementary Figure 2 and Supplementary Table 5). Latent noncognitive factors were also modestly correlated with latent factors of general cognitive ability (Supplementary Table 6) at the same age (Supplementary Table 7).

We examined whether general cognitive ability could account for the associations between noncognitive skills and academic achievement. Multiple regression analyses showed that both noncognitive factors were substantially and significantly associated with academic achievement beyond cognitive skills at every stage of compulsory education (Figure 2A and Supplementary Table 8). The relative contribution of noncognitive skills to academic achievement increased developmentally, particularly when considering self-reported measures. For self-reported education-specific noncognitive skills, the effect size of the relative prediction of achievement increased from *β* = .10 (SE = 0.02) at age 9 (effect size for cognitive ability: *β* = .46, SE = 0.01) to *β* = .28 (SE = 0.02) at age 12 (effect size for cognitive ability: *β* = .36, SE = 0.02) to *β* = .58 (SE = 0.02) at age 16 (effect size for cognitive ability: *β* = .39, SE = 0.01). A developmental increase was also observed for self-reported measures of domain-general self-regulation skills, for which the predictive power increased from *β* = .11 (SE = 0.02) at age 9 to *β*= .21 (SE = 0.01) at age 16, after accounting for general cognitive ability (Supplementary Table 8).

**Figure 2.**
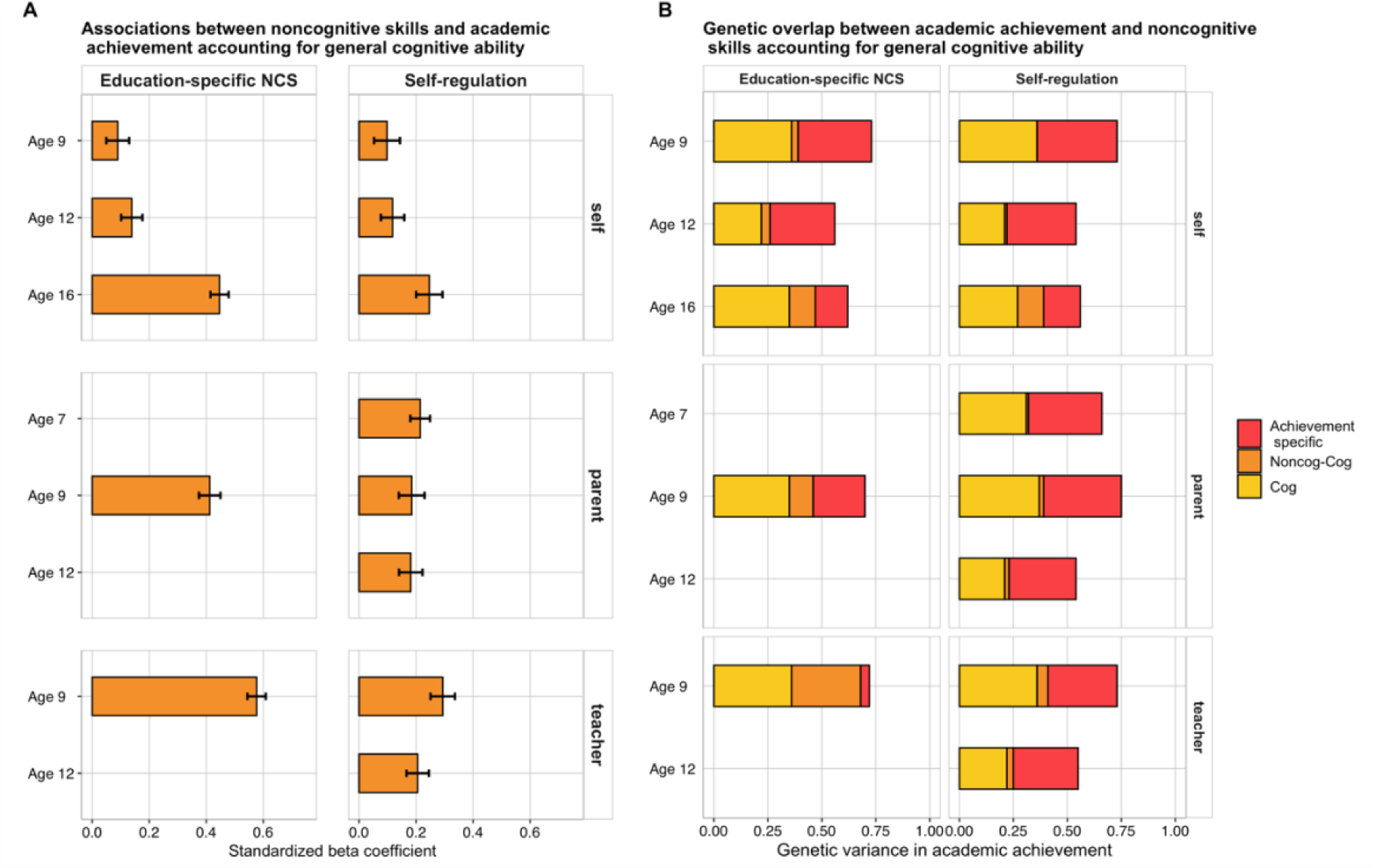
Associations between noncognitive skills and academic achievement accounting for general cognitive ability. **(Panel A)**. Associations between latent factors of noncognitive skills and academic achievement at ages 7, 9, 12 and 16, after accounting for general cognitive ability at the same age using multiple regression. Each bar indicates the effect size of standardized regression coefficients, and the error bars indicate the 95% confidence intervals around the estimates. The left panel shows the associations for latent measures of education-specific noncognitive skills (NCS), while the right panel the associations for latent dimensions of domain-general self-regulation skills. The figure is further divided into self-rated (top panel), parent-rated (middle panel) and teacher-rated (bottom panel) measures. **(Panel B)**. Each bar represents genetic effects (standardized and squared path estimates) on academic achievement over development and includes three shadings. The lighter (yellow) shadings indicate the proportion of genetic variance in academic achievement that can be attributed to genetic variance in cognitive skills (Cog). The orange shadings indicate the proportion of genetic variance in academic achievement that can be attributed to genetic variance in noncognitive skills, independent of the genetics of cognitive skills (Noncog – Cog). The red shadings indicate genetic effects on academic achievement independent of the genetics of cognitive and noncognitive skills (Achievement specific). Results are further divided into self-rated (top panel), parent-rated (middle panel) and teacher-rated (bottom panel) measures. Standardized paths and 95% Confidence intervals for all estimates are presented in Supplementary Tables 12 and 13.

### Specific genetic associations between noncognitive skills and academic achievement persist after accounting for cognitive skills and increase in magnitude across development

Applying twin designs (Methods), we found that the heritability (i.e., the extent to which observed differences in a trait are accounted for by genetic differences) of noncognitive skills differed significantly across raters and developmental stages (Supplementary Note 2, Supplementary Table 9, and Supplementary Figures 3-7). The heritabilities of latent noncognitive factors, which exclude error of measurement, ranged between 70% (95% CIs = 0.63; 0.77) for self-reported education-specific skills at age 9 and 93% (95% CIs = 0.91; 0.96) for parent-reported education-specific noncognitive skills at age 9 (Supplementary Note 2, Supplementary Tables 10-11 and Supplementary Figure 8). These substantial heritability estimates are consistent with previous studies that investigated the heritability of latent dimensions of noncognitive skills^11^ and of a general factor of psychopathology across different raters^40^. The correlation between noncognitive measures and academic achievement was mostly accounted for by genetic factors and, to a lesser extent, by nonshared environmental factors (Supplementary Note 2 and Supplementary Figure 8).

We then investigated whether the observed genetic associations between latent noncognitive factors and academic achievement could be accounted for by genetic factors associated with cognitive skills. We investigated this question with a series of trivariate Cholesky decompositions (Methods) the results of which are presented in Figure 2B, which reports standardized squared path estimates, and Supplementary Tables 12 and 13, which report standardized path estimates and 95% confidence intervals. The Cholesky approach, similar to hierarchical regression, parses the genetic and environmental variation in each trait into that which is accounted for by traits that have been previously entered into the model and the variance which is unique to a newly entered trait.

Each bar in Figure 2B is the outcome of a different trivariate Cholesky decomposition of the heritability of academic achievement (the total length of the bar) into genetic effects associated with noncognitive skills after controlling for genetic effects associated with cognitive skills at the same age. We found that genetic effects associated with cognitive skills accounted for between 21% and 36% of the total variance in academic achievement, as indicated by standardized paths ranging between 0.46 (95% CIs = 0.37; 0.54) and 0.60 (95% CIs = 0.50; 0.70). Genetic effects associated with noncognitive skills, independent of cognitive skills, accounted for between 0.1% and 32.5% of the variance in academic achievement, independent of cognitive skills, standardized paths ranged between 0.01 (95% CIs = −0.16; 0.17) for self-reported self-regulation at age 9 and 0.57 (95%CIs = 0.48; 0.67) for teacher reported education-specific noncognitive skills at age 9. Lastly, we found that between 5% and 37% of the variance in academic achievement was independent of genetic effects associated with cognitive and noncognitive skills; Standardized paths ranged between 0.23 (95%Cis = 0.13; 0.33) and 0.61 (0.52; 0.70).

The top three rows of Figure 2B illustrate the developmental increase in how the genetics of self-reported noncognitive skills contribute to the genetics of academic achievement. Focusing on education-specific noncognitive skills, we found that standardized squared path estimates increased from 1% of the total variance in academic achievement at age 9 (standardized path estimate = 0.01 [95% CIs = −0.16; 0.17]) to 4% at age 12 (standardized path estimate = 0.16 [95% CIs = 0.02; 0.30]) and 12% at age 16 (standardized path estimate = 0.35 [95% CIs = 0.26; 0.44]) (Supplementary Tables 12 and 13). This increased contribution beyond cognitive skills was also observed for domain-general self-regulation. See Supplementary Figure 9 for the full models’ results which include shared and nonshared environmental estimates.

### A new PGS of noncognitive skills calculated by extending the GWAS-by-subtraction approach

In order to obtain a PGS for use in subsequent analyses, we first extended previous work using the GWAS-by-subtraction approach to identify genetic variants associated with non-cognitive skills.^20^ Previous GWAS-by-subtraction work leveraged Genomic structural equation modelling (SEM)^41^ and the two genome-wide association studies (GWAS) of educational attainment and cognitive performance to separate the genetic variance in educational attainment into a cognitive component and a residual, noncognitive component. We extended this model in two directions. First, we extended the latent cognitive factor by including GWAS summary statistics from additional cognitive measures (episodic memory; processing speed, executive functions, and reaction time).^42^ Second, we included other socioeconomic attainment variables, including Townsend Deprivation and Income,^43^ in addition to educational attainment^17^. The resulting noncognitive factor can therefore be defined as genetic variation shared by educational attainment, income, and neighborhood deprivation that is independent of all measured cognitive abilities. Akin to Demange et al. 2021, we then fitted a Cholesky model (Methods) where indicators of the noncognitive latent factor (henceforth NonCog) were regressed on the cognitive latent factor (henceforth Cog; Figure 3A and Supplementary Table 14).

**Figure 3.**
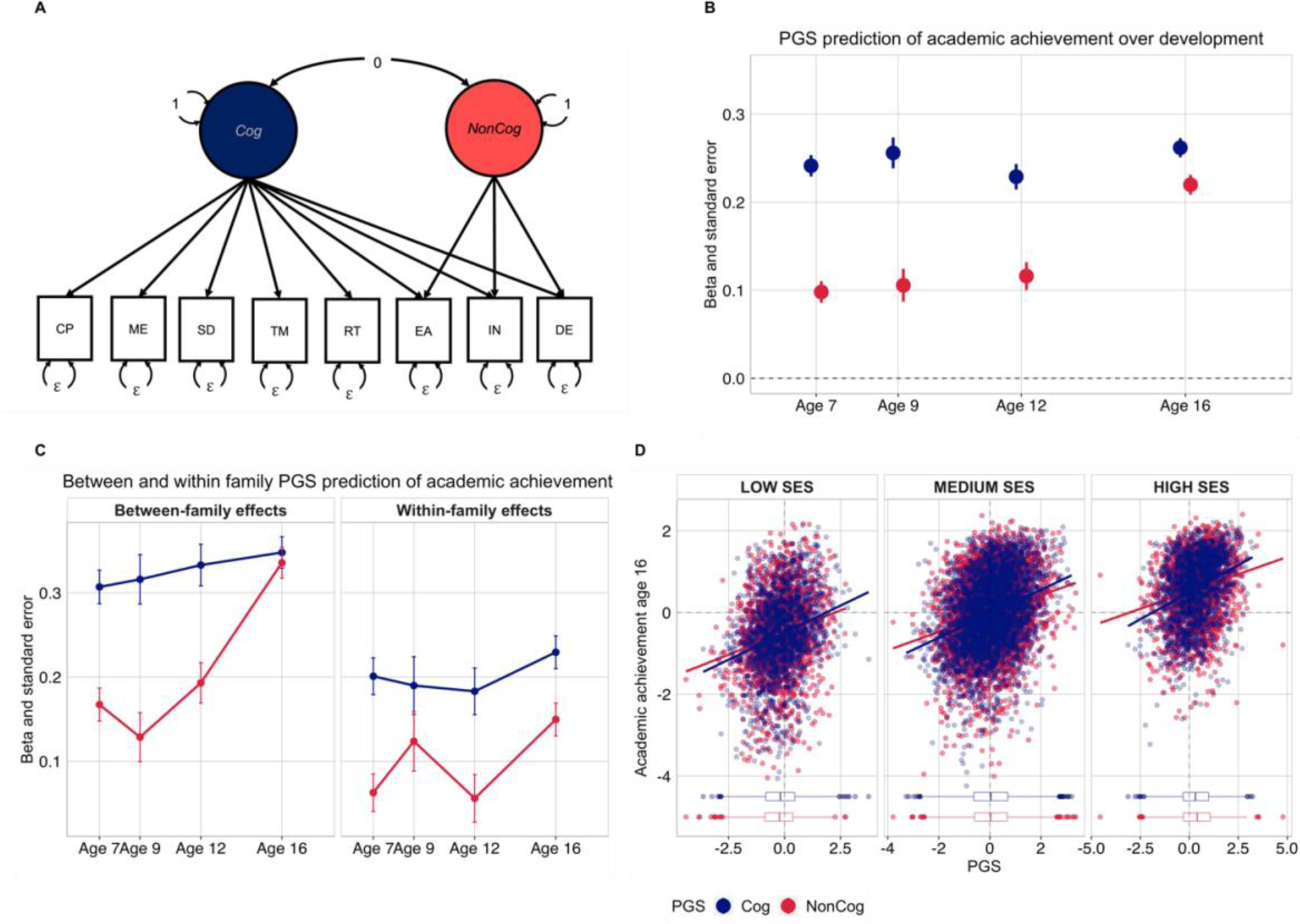
Contribution of noncognitive genetics to academic development: genomic analyses and gene-environment interplay. (Panel A.) Path diagram for the extension of the GWAS-by-subtraction model implemented in genomic structural equation model. In addition, GWAS summary statistics for cognitive performance (CP) and educational attainment (EA), summary statistics of memory (ME), symbol digit (SD), trail making (TM), and reaction time (RT) GWASs loaded on the cognitive (Cog) latent factor while GWAS summary statistics for income (IN) and deprivation (DE) loaded on the noncognitive (NonCog) latent factor, in addition to EA (Methods). (Panel B.) Cognitive and noncognitive polygenic score (PGS) prediction of academic achievement at ages 7,9, 12 and 16. (Panel C.) Results of polygenic scores analyses after partitioning the effects of Cog and NonCog into between and within family factors. (Panel D.) Cognitive (Cog) and noncognitive (Noncog) PGS prediction of academic achievement at the end of compulsory education (age 16), plotted at different levels of family socioeconomic status (SES).

The newly-created cognitive and noncognitive factors correlated strongly with those obtained from Demange et al.^20^ (Supplementary Table 15). The genetic correlation was 0.96 for the cognitive factors and 0.93 for the noncognitive factors. The genetic correlation between Cog and NonCog was rg = 0.15. Supplementary Figure 10 shows the genetic correlations between the newly created Cog and NonCog genetics and 18 psychiatric, personality and socio-economic traits, which we compared to the genetic correlations obtained by Demange et al.^20^. The pattern of associations was largely consistent across the two models. However, in some instances, results diverged. Specifically, with respect to psychiatric traits, autism, anorexia, and ADHD, a larger gap was observed between the cognitive and noncognitive factors, as compared to Demange et al., where differences in the correlations were less pronounced or absent. As expected, the results differed most for socioeconomic traits, with stronger correlations for NonCog than Cog with longevity (r = 0.52, SE = 0.04, p = 1.04E-45 Vs. r = 0.35, SE =0.03 p = 6.40E-31), neighbourhood deprivation (r = −0.66, SE = 0.04, p = 3.85E-54 Vs. r = −0.28, SE = 0.04, p = 5.98E-12), and educational attainment (r = 0.83, SE = 0.01, p = 0.00E+00 Vs. r = 0.65, SE = 0.01, p = 0.00E+00; Supplementary Figure 10 and Supplementary Table 15).

### The noncognitive polygenic prediction of academic achievement increases over development

We calculated polygenic scores (PGS) for Cog and NonCog and examined their association with cognitive, noncognitive and academic phenotypes over development. Polygenic scores leverage findings from GWAS and aggregate single-nucleotide polymorphisms (SNPs) across the genome into a single composite index summarizing genetic influence on a target trait. We calculated PGS as the sum of SNPs at all loci weighted by the effect size of their association (see Methods). We first investigated whether and to what extent Cog and NonCog PGS predicted individual differences in noncognitive skills across development by modelling both PGSs in a multiple regression model (Methods). In line with our previously obtained results showing a moderate association between cognitive and noncognitive traits, we found that the Cog PGS significantly predicted variation in noncognitive skills across development, with standardized effect sizes ranging between ß = 0.04, SE = 0.02 and ß = 0.22, SE = 0.02 (Figure S11 and Supplementary Table 16). The NonCog PGS, independent of the cognitive PGS, predicted observed variation in noncognitive skills at all developmental stages. Associations were small at earlier ages (e.g., ß = 0.07, SE = 0.02, p(corrected) = 1.93E-03) for parent-reported education-specific noncognitive skills at 9, and ß = 0.10, SE = 0.01, p(corrected) = 2.24E-11 for parent-reported self-regulation at 7) but they increased developmentally, particularly for self-reported education-specific noncognitive measures (ß = 0.16, SE = 0.02, p(corrected) = 8.30E-17 at age 16). The only exception was observed for self-reported education-specific noncognitive skills at age 9, for which the prediction was negative (ß = −0.03, SE = 0.02) and did not reach significance after accounting for multiple testing (Supplementary Table 16). In Supplementary Note 3a, we show that this increase in prediction was significant overtime for the NonCog PGS, but not for the Cog PGS. Furthermore, we show that this increase is not explained by the NonCog PGS capturing more cognitive variance later in adolescence (Supplementary note 3b), or by socio-economic status (Supplementary note 3c).

Cog and NonCog PGSs predicted variation in general cognitive ability, verbal ability, and nonverbal ability at all developmental stages. As expected, the Cog PGS prediction of cognitive phenotypes was substantially stronger than the NonCog prediction, with estimates ranging between ß = 0.19, SE = 0.1, p(corrected) = 3.77E-42and ß = 0.27, SE = 0.02, p(corrected) = 1.04E-52 for the Cog PGS and between ß = 0.10,SE = 0.02 p(corrected) = 4.41E-10 and ß = 0.18, SE = 0.02, p(corrected) 5.51E-21 for the NonCog PGS (Supplementary Table 16).

Next, we considered the effects of the Cog and NonCog PGSs on academic achievement over development. We detected associations between the Cog PGS and achievement as early as age 7 (ß =0.24, SE = 0.01, p(corrected) = 3.68E-86), these associations remained largely consistent across development (ß = 0.26, se = 0.01, p(corrected) = 2.71E-126 at age 16). Although we observed weaker effects for the NonCog PGS in early childhood (ß =0.10, SE = 0.01, p(corrected) = 8.12E-15) as compared to the Cog PGS, these increased across development and reached effects comparable to those of the Cog PGS at age 16 (ß =0.22, SE = 0.01, p(corrected) = 1.85E-84; Figure 3B and Supplementary Table 16). The same pattern of associations was observed also when considering achievement in English and mathematics, separately (Supplementary Table 16). This observed increase in the NonCog PGS prediction of academic achievement over development is consistent with transactional models of gene-environment correlation, driven by noncognitive genetics. These PGS predictions were in line with those obtained from the PGSs created using the GWAS-by-subtraction method published by Demange et al. (Supplementary Table 17).

### Differences between siblings in the polygenic prediction of academic achievement across development

Given our observation of an increase in the NonCog PGS contributions to academic achievement across development, we extended our preregistered analyses (https://osf.io/m5f7j/) to examine whether and to what extent this increase was accounted for by family-wide processes. Specifically, using a sibling difference design we separated the NonCog PGS contributions into within-family effects, indexing direct genetic effects, from between-family effects, which may include indirect genetic effects and demographic confounding (Methods). We examined within and between family contributions of the Cog and NonCog PGS on academic achievement from age 7 to 16.

Two main findings emerged from this analysis (Figure 3C). First, we observed that the effect sizes for the direct effects of NonCog were about half as the size of the population-level associations (Supplementary Table 18). Similarly, the prediction from the Cog PGS was reduced by over one-third, consistent with previous evidence^44^. Second, while the Cog direct and indirect genetic effects did not vary substantially over the developmental period considered (from ß = 0.20, SE = 0.02, p = 2.75E-20 to ß = 0.23, SE = 0.02, p = 4.12E-32), NonCog effects showed an increase from age 7 to age 16 (from ß = 0.06, SE = 0.02, p = 0.005 to ß = 0.15, SE = 0.02, p = 1.39E-14; Figure 3C, and Supplementary Table 18). These results suggested that the developmental increase in the between family PGS prediction was mostly driven by noncognitive rather than cognitive skills. In addition, this developmental increase could be observed for both indirect and direct genetic effects. We conducted sensitivity analyses, and replicated the results, with the PGSs constructed using the method published by Demange et al. (Supplementary Table 18b).

### Does socioeconomic status modify the association between Cog/NonCog PGS and educational outcomes across development?

Lastly, we extended our preregistered analyses to test whether socio-economic status (SES) could explain or modify the observed pattern of developmental associations between PGS and academic achievement. We fitted multivariable models at each developmental stage including Cog/NonCog PGS effects, along with SES at recruitment, covariates, and their two-way interactions (see Methods) to test whether SES moderated Cog and NonCog PGS effects on academic achievement. After adjusting for SES, the same pattern of relationships was observed, with a relatively stable association between the Cog PGS and achievement, and a steeper increase in the NonCog PGS prediction, even though all effects were attenuated (Supplementary Table 19). We did not detect significant interaction effects between either the Cog or the NonCog PGS with SES (Supplementary Table 19).

Figure 3D depicts mutually adjusted slopes for the Cog and NonCog PGS prediction against academic achievement at different levels of family SES. The figure shows that although higher SES corresponded to greater achievement on average, the slope of the association between the Cog and NonCog PGS and achievement did not differ across socio-economic strata. Higher PGS, for both cognitive and noncognitive skills, corresponded to higher academic achievement, and higher SES corresponded to both higher mean PGSs and higher achievement, indicating a correlation rather than an interaction between genetic and environmental influences on academic achievement.

## Discussion

We investigated the contribution of cognitive and noncognitive genetics to academic achievement during compulsory education in a UK-based sample. Four complementary findings emerged. First, noncognitive skills increasingly predicted academic achievement over the school years, and these effects remain substantial even after accounting for cognitive skills. Second, the contribution of noncognitive skills to academic achievement is mainly due to common genetic factors, whose influence also increases over the school years. For example, the noncognitive polygenic score prediction of academic achievement nearly doubles over the school years, while the cognitive polygenic score prediction remains relatively stable. Third, the increasingly important role of noncognitive genetics persists even after accounting for family-fixed effects. Fourth, polygenic score contributions to academic development did not differ across socio-economic contexts. Together, these findings highlight the important role that noncognitive skills play during primary and secondary education and suggest that fostering such skills might provide an avenue for successful educational strategies and interventions.

The first set of novel findings about development emerged from twin analyses of the covariance between noncognitive traits and academic achievement. First, we found that genetic factors accounted for most of the observed correlations between noncognitive skills and academic achievement at all developmental stages. Second, both phenotypic and genetic correlations increased developmentally, particularly for self-reported measures of noncognitive traits. Third, our twin analyses showed that genetic factors accounted for most of the correlations between noncognitive skills and academic achievement after accounting for cognitive skills. Finally, this independent genetic contribution of noncognitive skills to academic achievement increased developmentally. This increase was observed for both education-specific noncognitive skills, where the measures included in the general factors changed developmentally, as well as for domain-general self-regulation skills, for which the same measures were collected at all developmental stages. Therefore, the observed developmental increase in phenotypic and genetic associations independent of cognitive skills is unlikely to be an artefact of inconsistencies in measurement but rather reflects the increasingly important role of noncognitive skills across compulsory education.

A further aim of the current study was to better understand what was captured by the noncognitive PGS constructed using GWAS-by-subtraction^20^, particularly in relation to what other skills beyond cognitive ability propel students down different educational trajectories. Given the link between socioeconomic status and academic achievement^45^ we were specifically interested in whether the noncognitive PGS also indexed socio-economic-related factors. To this end, we extended the GWAS-by-subtraction model in two directions. First, with the aim of making a more refined cognitive factor, we added summary statistics from several other GWASs of fluid intelligence. Second, we included GWASs of other traits known to associate with achievement beyond cognitive abilities, specifically targeting SES-related traits such as income and social deprivation, making the noncognitive PGS factor more explicitly socio-economic relevant.

It should be highlighted that GWAS of SES-relevant measures may be more subject to sociodemographic confounds such that estimates of SNP effects will also capture population stratification phenomena, such as geographic clustering^46^. This limitation is particularly relevant for the GWAS of social deprivation, as the measure is an area-based score of social inequality. Interestingly, the results obtained from this new model paralleled those we obtained when we applied the cognitive and noncognitive PGSs from the original GWAS-by-subtraction model, which only used educational attainment to define the noncognitive factor. This suggests that the PGS measure of noncognitive skills from Demange et al. may have already captured some SES-related effects. Importantly, our employment of a within-family comparison helped us to mitigate possible confounds associated with uncontrolled population stratification.

Paralleling our multivariate twin results, we observed that the effects of the prediction from noncognitive PGS to academic achievement increased from childhood to adolescence, beyond the effects of the cognitive PGS. A few explanations are possible for this finding. First, this could be attributable to gene-environment correlation (rGE), which could be passive, evocative, or active^47,48^. Another explanation could be that PGS become increasingly predictive during development because the sample becomes closer in age to the adult samples where GWAS effect sizes were estimated in the case of educational attainment and cognitive performance^17^. However, it is of note that this increase in prediction was not observed for the cognitive PGS, for which effects on academic achievement were mostly stable developmentally. Moreover, our triangulation of results across multiple methods (including phenotypic and twin analyses) adds support to our finding of these developmental differences between cognitive and noncognitive genetics.

We applied a within-sibling design^44^ to test whether environmental variables that are shared by siblings and that potentially confound PGS could explain the observed increase in the predictive power of the noncognitive PGS. While the contributions of both PGSs were attenuated within-family, suggesting a substantial role for environmental confounds shared by family members, an increase in the contribution of noncognitive PGS to academic achievement from age 7 to 16 was still evident when comparing siblings. In contrast, the within-family contribution of the cognitive PGS remained relatively stable. The increase in the noncognitive PGS prediction at the within-family level is consistent with transactional processes driven by active or evocative gene-environment correlation^31,48,49^ for noncognitive PGS. As children grow up, they actively evoke or shape their environmental experiences based in part on their genetic dispositions, and these experiences in turn contribute to their academic development. Our findings suggest that children’s educational experiences are increasingly shaped by their propensity towards noncognitive skills.

To delve deeper into the role of socio-economic factors, we tested whether SES could modify the relationship between cognitive and noncognitive PGSs and academic achievement over development. While we did not find evidence for interaction effects in this regard, the cognitive and noncognitive PGS were conditionally independent in a multivariable model including SES, further indicating that the genetics captured by the noncognitive skills factor was at least partly independent of SES-related genetic and environmental effects.

One caveat of these gene-environment interaction analyses is that adjusting for a heritable covariate, like SES, can yield biased estimates in multivariable models including PGS^50,51^. Future work is needed to determine whether this is the case, perhaps leveraging results of within-family GWAS to construct PGS for ‘direct’ effects within families^52^. This limitation also pertains to our within-sibling PGS analyses, as it might be difficult to separate direct and indirect effects using population-based GWAS effects as a starting point^53^. Follow-up of these analyses employing PGS for direct effects obtained from family-based GWAS will shed light on this potential limitation. A further caveat of the present work is that, while we investigated genetic effects on noncognitive skills and their link with academic achievement across development, we did not investigate stability and change using longitudinal models. Future work explicitly investigating developmental change at the phenotypic^54^, genetic^29^ and genomic^41,55^ level, for example using latent growth models^56^, will address further developmental questions related to the role of noncognitive skills in academic development.

To conclude, our study provides an in-depth investigation of the role of noncognitive genetics in academic development. Triangulating multiple genetic and genomic methods, we found consistent evidence for the increasingly important role that noncognitive skills play during compulsory education. Genetic dispositions towards noncognitive skills become increasingly predictive of academic achievement and, by late adolescence, they explain as much variance in achievement as do genetic dispositions towards cognitive skills. Results from within-family and developmental analyses are consistent with theorized transactional processes of active/evocative gene-environment correlation by which, as they grow up, children evoke and actively select academic environments that correlate with their genetic disposition towards noncognitive skills^47,48^. Fostering noncognitive skills might provide a successful avenue for educational interventions.

## Methods

### Sample

Participants are part of the Twins Early Development Study (TEDS), a longitudinal study of twins born in England and Wales between 1994 and 1996. The families in TEDS are representative of the British population for their cohort in terms of socio-economic distribution, ethnicity and parental occupation. Ten thousand families are still actively involved with the TEDS study over twenty years after the first data collection wave (see^57^ for additional information on the TEDS sample). The present study includes data collected in TEDS across multiple waves. Specifically, we will analyze data collected over five collection waves, when the twins were 4, 7, 9, 12 and 16 years old. The sample size differs between collection waves, numbers for all measures included in the study are reported in Supplementary Table 1.

### Measures

Below we provide a brief description of all the measures included in the present study. Please refer to https://www.teds.ac.uk/datadictionary for detailed descriptions of each measure and information on the items included in each construct.

#### Education-specific noncognitive skills

At **age 9** data on education-specific noncognitive skills were collected from parents, teachers and self-reports from the twins. Measures of academic self-perceived ability^58^, academic interest^58^ and the Classroom Environment Questionnaire (CEQ^59^) were available from all raters. The CEQ included the following subscales rated by parents and twins: (1) CEQ classroom satisfaction scale; (2) CEQ educational opportunities scale; (3) CEQ adventures scales, assessing enjoyment of learning. Ratings on the CEQ classroom satisfaction scale were also provided by the teachers.

At **age 12** data on education-specific noncognitive skills were collected from parents, teachers, and self-reports. The following measures were collected: academic self-perceived ability^58^, academic interest^58^, the mathematics environment questionnaire^60^ and the literacy environment questionnaire^61^. The questionnaires asked several questions related to literacy and mathematics, including items such as: *Reading is one of my favourite activities; When I read books, I learn a lot*; and *In school, how often do you do maths problems from text books?* all rated on a four-point Likert scale.

At **age 16** education-specific noncognitive skills were assessed via self-reports provided by the twins. The battery of education-specific noncognitive constructs included the following measures:

a. The <u>brief academic self-concept</u> scale included 10 items (adapted from^62^), such as: *I like having difficult work to do* and *I am clever*, rated on a 5-point Likert scale.
b. <u>School engagement</u>^63^ includes 5 subscales: teacher-student relations; control and relevance of schoolwork; peer support for learning; future aspirations and goals; family support for learning. The school engagement scale includes items such as: *I enjoy talking to the teachers at my school*, *I feel like I have a say about what happens to me at school*, *School is important for achieving my future goals*, and *When I have problems at school, my family/carer(s) are willing to help me*, rated on a 4-point Likert scale.
c. <u>Grit</u> was assessed with 8 items from the Short Grit Scale (GRIT-S)^64^ asking the twins to report on their academic perseverance answering questions such as: *Setbacks don’t discourage me*, and *I am a hard worker,* rated on a 5-point Likert scale.
d. <u>Academic ambition</u>^65^ was measured with 5 items asking participants to rate statements like the following on a 5-point Likert scale: *I am ambitious* and *achieving something of lasting importance is the highest goal in life*.
e. <u>Time spent studying mathematics</u> was assessed with 3 items asking participants how much time every week they spent in: *Regular lessons in mathematics at school, Out-of school-time lessons in mathematics*, and *Study or homework in mathematics by themselves*.
f. <u>Mathematics self-efficacy</u>^66^ was measured with 8 items asking students how confident they felt about having to perform different mathematics tasks, for example: *Calculating how many square metres of tiles you need to cover a floor* and *Understanding graphs presented in newspapers*, rated on a 4-point Likert scale
g. <u>Mathematics interest</u>^66^ asked participants to respond to 3 questions related to interest in mathematics, including: I *do mathematics because I enjoy it* and *I am interested in the things I learn in mathematics*.
h. <u>Curiosity</u> was assessed with 7 items^67^ asking participants to rate statements such as: *When I am actively interested in something, it takes a great deal to interrupt me* and *Everywhere I go, I am looking out for new things or experiences* on a 7-point Likert scale
i. <u>Attitudes towards school</u> was measured using the PISA attitudes to school measure^66^ which included 4 items such as: *School has helped give me confidence to make decisions* and *School has taught me things which could be useful in a job* rated on a 4-point Likert scale.

#### Self-regulation

Emotional and behavioral self-regulation was assessed at all ages using the Strengths and Difficulties Questionnaire (SDQ)^68^. Data on domain-general self-regulation skills was collected from parents, teachers and self-reported by the twins. The SDQ includes 5 subscales: hyperactivity, conduct problems, peer problems, emotional problems, and prosocial behaviour. Composite scores for all subscales except prosocial behaviour were reversed so that higher scores indicated higher levels of domain-general self-regulation skills. At age 7, domain-general self-regulation skills were rated by the parents; at age 9 and 12 by the parents, teachers and self-reported by the twins; and at age 16 self-reported by the twins.

#### Cognitive ability

At **age 7** cognitive ability was measured using four cognitive tests that were administered over the telephone by trained research assistants. Two tests assessed verbal cognitive ability: a 13-item Similarity test and 18-item Vocabulary test, both derived from the Wechsler Intelligence Scale for Children (WISC-III)^69^. Nonverbal cognitive ability was measured using two tests: a 9-item Conceptual Groupings Test^70^, and a 21-item WISC Picture Completion Test^69^. Verbal and nonverbal ability composites were created taking the mean of the standardized test scores within each domain. A *g* composite was derived taking the mean of the two standardized verbal and two standardized nonverbal test scores.

At **age 9** cognitive ability was assessed using four cognitive tests that were administered as booklets sent to TEDS families by post. Verbal ability was measured using the first 20 items from WISC-III-PI Words test^71^ and the first 18 items from WISC-III-PI General Knowledge test^71^. Nonverbal ability was assessed using the Shapes test (CAT3 Figure Classification)^72^ and the Puzzle test (CAT3 Figure Analogies)^72^. Verbal and nonverbal ability composites were created taking the mean of the standardized test scores within each domain. A *g* composite was derived taking the mean of the two standardized verbal and two standardized nonverbal test scores.

At **age 12**, cognitive ability was measured using four cognitive tests that were administered online. Verbal ability was measured using the full versions of the verbal ability tests administered at age 9: the full 30 items from WISC-III-PI Words test^71^ and 30 items from WISC-III-PI General Knowledge test^71^. Nonverbal ability was measured with the 24-item Pattern test (derived from the Raven’s Standard Progressive Matrices)^73^ and the 30-item Picture Completion test (WISC-III-UK)^69^. Verbal and nonverbal ability composites were created taking the mean of the standardized test scores within each domain. A *g* composite was derived from the mean of the two standardized verbal and two standardized nonverbal test scores.

At **age 16** cognitive ability was assessed using a composite of one verbal and one nonverbal test administered online. Verbal ability was assessed using an adaptation of the Mill Hill Vocabulary test^74^, Nonverbal ability was measured using an adapted version of the Raven’s Standard Progressive Matrices test^73^. A *g* composite was derived taking the mean of the two standardized tests.

#### Academic achievement

At **age 7** academic achievement was measured with standardized teacher reports and consisted of standardized mean scores of students’ achievements in English and mathematics, in line with the National Curriculum Level. Performance in English was assessed in four domains: speaking, listening, reading, and writing abilities; performance in maths was assessed in three domains: applying mathematics, as well as knowledge about numbers, shapes, space and measures.

At **age 9,** academic achievement was again assessed using teacher reports. The domains assessed were the same for English and mathematics (although on age-appropriate content). In addition, performance in science was assessed considering two key domains: scientific enquiry and knowledge and understanding of life processes, living things and physical processes.

At **age 12**, academic achievement was assessed in the same way as at age 9, with two exceptions. Mathematics added a fourth domain, data handling, and science added a third domain, materials and their properties. These additions were in line with the changes made to the National Curriculum teacher ratings.

At **age 16**, academic achievement was measured using the General Certificate of Secondary Education (GCSE) exam scores. The GCSE is the UK nationwide examination usually taken by 16-year-olds at the end of compulsory secondary education^75^. Twins’ GCSE scores were obtained via mailing examination results forms to the families shortly after completion of the GCSE exams by the twins. For the GCSE, students could choose from a wide range of subjects. In the current analyses the mean score of the three compulsory GCSE subjects: English Language and/or English Literature, mathematics, and a science composite (a mean score of any of the scientific subjects taken, including physics, chemistry, and biology).

#### Family socio-economic status

At first contact, parents of TEDS twins received a questionnaire by post, and were asked to provide information about their educational qualifications, employment, and mothers’ age at first birth. A socioeconomic status composite was created by standardizing these three variables and calculating their mean. The same measures, except for mother’s age at first birth, were used to measure family socioeconomic status at age 7. At age 16, data on socioeconomic status were collected using a web questionnaire, and a total score was calculated from the standardized mean of 5 items: household income, mother’s and father’s highest qualifications, and mother’s and father’s employment status.

#### Genetic data

Two different genotyping platforms were used because genotyping was undertaken in two separate waves, 5 years apart. AffymetrixGeneChip 6.0 SNP arrays were used to genotype 3,665 individuals. Additionally, 8,122 individuals (including 3,607 DZ co-twin samples) were genotyped on Illumina HumanOmniExpressExome-8v1.2 arrays. Genotypes from a total of 10,346 samples (including 3,320 DZ twin pairs and 7,026 unrelated individuals) passed quality control, including 3,057 individuals genotyped on Affymetrix and 7,289 individuals genotyped on Illumina. The final data contained 7,363,646 genotyped or well-imputed SNPs. For additional information on the treatment of these samples see^76^.

### Analytic strategies

#### Phenotypic analyses: Confirmatory factor analysis, correlations, and regressions

Confirmatory factor analysis (CFA) was employed to create latent dimensions of noncognitive skills and general cognitive ability at all ages. Based on the well-established literature on general cognitive ability (g) and previous work in the TEDS sample^77^, we constructed one factor for g at each developmental stage. Each g factor was created by taking the weighted loadings of two verbal and two nonverbal tests (see Measures and Supplementary Table 6). CFA was also employed to construct dimensions of noncognitive characteristics. Based on previous meta-analytic work on the noncognitive characteristics that matter for educational outcomes^9,78^, we embraced a theoretical distinction between education-specific noncognitive characteristics (e.g., motivations, attitudes and goals) and broader, more de-contextualized measures of self-regulation (e.g., behavioural and emotional regulation), and created separate factors for a) education-specific noncognitive characteristics and b) domain-general self-regulation skills separately for ages and raters, including all the measures available at each age for each rater (see Supplementary Tables 2 and 3 for factor loadings and model fit indices).

We applied phenotypic correlations to examine the associations between noncognitive skills (both observed measures and factors) and general cognitive ability and academic achievement at each age. We applied multiple regressions to explore the associations between noncognitive skills and academic achievement accounting for general cognitive ability. We applied Benjamini-Hochberg correction^79^ to account for multiple testing.

#### Genetic analyses: The twin method

The twin method allows for the decomposition of individual differences in a trait into genetic and environmental sources of variance by capitalizing on the genetic relatedness between monozygotic twins (MZ), who share 100% of their genetic makeup, and dizygotic twins (DZ), who share on average 50% of the genes that differ between individuals. The method is further grounded in the assumption that both types of twins who are raised in the same family share their rearing environments to approximately the same extent ^80^. By comparing how similar MZ and DZ twins are for a given trait (intraclass correlations), it is possible to estimate the relative contribution of genetic factors and environments to variation in that trait. Heritability, the amount of variance in a trait that can be attributed to genetic variance (A), can be roughly estimated as double the difference between the MZ and DZ twin intraclass correlations^80^. The ACE model further partitions the variance into shared environment (C), which describes the extent to which twins raised in the same family resemble each other beyond their shared genetic variance, and non-shared environment (E), which describes environmental variance that does not contribute to similarities between twin pairs (and also includes measurement error).

The twin method can be extended to the exploration of the covariance between two or more traits (multivariate genetic analysis). Multivariate genetic analysis allows for the decomposition of the covariance between multiple traits into genetic and environmental sources of variance, by modelling the cross-twin cross-trait covariances. Cross-twin cross-trait covariances describe the association between two variables, with twin 1’s score on variable 1 correlated with twin 2’s score on variable 2, which are calculated separately for MZ and DZ twins. The examination of shared variance between traits can be further extended to test the aetiology of the variance that is common between traits and of the residual variance that is specific to individual traits.

It is possible to apply structural equation modelling to decompose latent factors into A, C and E components, applying models such as the common pathway model. The **<u>common pathway model</u>** is a multivariate genetic model in which the variance common to all measures included in the analysis can be reduced to a common latent factor, for which the A, C and E components are estimated. As well as estimating the aetiology of the common latent factor, the model allows for the estimation of the A, C and E components of the residual variance in each measure that is not captured by the latent construct^81^.

A further multivariate twin method, grounded in SEM is the **<u>Cholesky decomposition</u>**, which examines the genetic and environmental underpinnings of the associations between multiple variables or latent factors. The Cholesky approach parses the genetic and environmental variation in each trait into that which is accounted for by traits that have been previously entered into the model and the variance which is unique to a newly entered trait. In our case the Cholesky decomposition partitions the genetic and environmental variance that is common across cognitive, noncognitive and achievement measures from the genetic and environmental variance that is common between noncognitive skills and achievement, independently of that accounted for by cognitive ability. Cholesky decompositions were conducted on latent dimensions of cognitive and noncognitive skills and observed variation in academic achievement (see Supplementary Tables 12 and 13).

#### Genetic analyses: Genomic structural equation model (SEM)

Genomic SEM^41^ is an approach to conduct multivariate genome-wide association (GWA) analyses. Based on the principles of SEM widely used in twin analyses and integrated with LD score regression^82^, Genomic SEM jointly analyzes GWA summary statistics for multiple traits to test hypotheses about the structure of the genetic covariance between traits. Here we employed Genomic SEM to create latent GWAS summary statistics for unmeasured traits based on other traits for which GWAS summary statistics exist. Recent work applied a GWAS-by-subtraction approach^20^ leveraging GWA studies of educational attainment (EA^17^) and cognitive performance (CP^17,83^) to obtain a GWA of noncognitive skills. The GWAS-by-subtraction approach estimates, for each single nucleotide polymorphism (SNP), an effect on EA that is independent of that SNP’s effect on CP (therefore indexing residual *noncognitive* SNP effects). The model regresses the EA and CP summary statistics on two latent variables, *Cog* and *NonCog*. EA and CP are both regressed on the *Cog* latent variable and only EA is regressed on the *NonCog* latent factor. The *Cog* and *NonCog* factors are specified to be uncorrelated and residual covariances across factor indicators are set to zero. *Cog* and *NonCog* are then regressed on each SNP, iterating across all SNPs in the genome.

We extended the GWAS-by-subtraction with the aim of obtaining potentially more fine-grained cognitive and noncognitive factors. Specifically, the model was extended as follows: Loading exclusively on the *Cog* factor: five UK Biobank cognitive traits (Cognitive Performance^83^, Symbol Digit Substitution, Memory, Trail Making Test and Reaction Time)^42^. Loading on both the *Cog* and *Noncog* factors: educational attainment^17^, Townsend deprivation index (http://www.nealelab.is/uk-biobank/), and income^43^. An additional difference from the original GWAS-by-subtraction is that we let residual variances vary freely (i.e., we did not constrain them to 0; see Figure 3A and Supplementary Table 14).

#### Genetic analyses: Construction of polygenic scores (PGS) and PGS analyses

Polygenic scores (PGS) were calculated as the weighted sums of each individual’s genotype across all single nucleotides polymorphisms (SNPs), using LDpred weights^84^. LDpred is a bayesian shrinkage method that corrects for local linkage disequilibrium (LD; i.e. correlations between SNPs) using information from a reference panel (we used the target sample (TEDS) limited to unrelated individuals) and a prior for the genetic architecture of the trait. We constructed PGS using an infinitesimal prior, that is assuming that all SNPs are involved in the genetic architecture of the trait, as this has been found to perform well with highly polygenic traits such as educational attainment, and in line with the approach adopted by Demange et al.^20^. In regression analyses, following from Demange et al.^20^, both the Cog and NonCog PGSs were included in multiple regressions together with the following covariates: age, sex, the first 10 principal components of ancestry, and genotyping chip and batch. We accounted for non-independence of observation using generalized estimating equation (GEE).

#### Genetic analyses: Within and between family analyses

We conducted within-sibling analyses using DZ twins to estimate family-fixed effects of both cog and non-cog PGS on achievement across development^44^. A mixed model was fit to the data including a random intercept to adjust for family clustering, and two family-fixed effects in addition to covariates (age, sex, the first 10 principal components of ancestry, and genotyping chip and batch): a between-family effect indexed by the mean family PGS (i.e., the average of the DZ twins’ PGS within a family), and a within-family effect, indexed by the difference between each twin’s PGS from the family mean PGS. Analyses were repeated with the PGS from Demange et al.^20^, as sensitivity analyses.

#### Genetic analyses: Gene x Environment interaction analyses

We conducted gene-environment (GxE) interaction analyses to test whether SES moderated the effects of the cognitive and noncognitive PGS prediction on academic achievement over development. We fit a linear mixed model including Cog and NonCog PGS (the extensions), SES and their two-way interactions after adjusting for covariates (as above) and two-way interactions between predictors and covariates, plus a random intercept to adjust for family clustering. We adjusted for multiple testing using the Benjamini–Hochberg false discovery rate (FDR) method^79^ for all PGS analyses, at an alpha level of .05.

## Acknowledgements

We gratefully acknowledge the ongoing contribution of the participants in the Twins Early Development Study (TEDS) and their families. TEDS has been supported by a program grant to RP from the UK Medical Research Council (MR/M021475/1 and previously G0901245), with additional support from the US National Institutes of Health (AG046938). MM is supported by a starting grant from the School of Biological and Behavioural Sciences at Queen Mary University of London. RP is supported by a Medical Research Council Professorship award (G19/2). KR is supported by a Sir Henry Wellcome Postdoctoral Fellowship. ADG was supported by NIH Grants R01MH120219 and RF1AG073593.

## Contributions

M.M., A.G.A. and R.P. conceived and designed the study.; M.M. and A.G.A. analyzed the data with helpful contributions from M.G.N. and P.B.; M.M., A.G.A., K.P.H., and R.P. wrote the paper with helpful contributions from M.G.N., P.B., K.R., R.C., S.v.S., P.D., E.v.B., A.G., L.R., J.d.F., J.B.P., E.M.T. All authors contributed to the interpretation of data, provided critical feedback on manuscript drafts, and approved the final draft.

## Code availability

Code will be available at https://github.com/CoDEresearchlab/NoncognitiveGenetics upon publication. Ahead of publication we will make the code available upon request.

