## Supplementary Notes and Figures for "Genetic contributions of noncognitive skills to academic development"

Malanchini, .M, Allegrini, A.G. et al.

### Supplementary Notes

**Note 1.** Associations between individual noncognitive measures, academic achievement, and general cognitive ability over development

**Note 2.** The aetiology of noncognitive measures and their association with academic achievement over development.

**Note 3.** Sensitivity Analyses.

### Supplementary Figures

**Figure 1.** Correlations between observed noncognitive measures and academic achievement from age 7 to 16.

**Figure 2**. Phenotypic associations between latent noncognitive measures, academic achievement, and general cognitive ability.

**Figure 3.** Heritability, shared environmental, and nonshared environmental estimates for all observed measures of education-specific noncognitive skills from age 9 to 16.

**Figure 4.** Heritability, shared environmental, and nonshared environmental estimates for all observed measures of low self-regulation from age 7 to 16.

**Figure 5.** Heritability, shared environmental, and nonshared environmental estimates for all observed measures of academic achievement from age 7 to 16.

**Figure 6.** Heritability, shared environmental, and nonshared environmental estimates for all observed measures of cognitive ability from age 7 to 16.

**Figure 7.** Heritability, shared environmental, and nonshared environmental estimates for all observed measures of early academic abilities at age 4.

**Figure 8.** Genetic and environmental influences on latent noncognitive dimensions and on their associations with academic achievement at the same age.

**Figure 9.** Contribution of genetic and environmental influences on cognitive and noncognitive skills to academic achievement at the same age.

**Figure 10.** Genetic correlations for the new cognitive and noncognitive factors obtained with the new extension of the GWAS-by-subtraction model.

**Figure 11.** Cognitive and noncognitive polygenic score (PGS) predictions of noncognitive, academic achievement and family socio-economic status phenotypes over development.

**Figure 12.** Cognitive and Noncognitive polygenic score (PGS) predictions of cognitive phenotypes over development.

### Supplementary Note 1

#### Associations between individual noncognitive measures and academic achievement over development

We examined the correlations between individual noncognitive traits and academic achievement at each age (Supplementary Figure 1 and Supplementary Table 1). Correlations between measures of education-specific noncognitive skills, including characteristics such as self-perceived ability, learning interest, enjoyment, and academic achievement were overall positive and ranged from *r* = 0.15 (95% CIs = 0.11, 0.19) for self-rated curiosity at age 16, to *r* = 0.81 (95% CIs =0.79, 0.82) for teacher-rated self-perceived ability at age 9 (Supplementary Figure 1A and Supplementary Table 1). Correlations between measures of self-regulation and educational achievement were modest at every developmental stage, with estimates ranging from very small effects, for example, *r* = 0.02 (95% CIs = -0.02, 0.05) for parent-rated prosocial behaviour at age 7 to moderate negative effect sizes, such as those observed for teacher-rated hyperactivity at age 9, *r* = -0.43 (95% CIs = -0.47, -0.41; see Supplementary Figure 1B and Supplementary Table 1).

When we considered latent dimensions of education-specific noncognitive skills and domain-general self-regulation skills, we found that individual differences in these factors of noncognitive characteristics were associated with variation in academic achievement at every stage during compulsory education, and associations became stronger developmentally. Because data on noncognitive characteristics were available from different raters (parents, teachers, and self-reports) we also created factors of noncognitive skills separately for each rater (Supplementary Tables 2 and 3). Effect sizes differed depending on the rater and developmental stage considered but tended to increase with age. For example, the association between self-rated education-specific noncognitive skills and academic achievement increased from small (*r* = 0.10) at age 9 to moderate (*r* = 0.41) at age 12, to strong (*r* = 0.51) at age 16. When considering parent and teacher ratings, we found that correlations between noncognitive factors and academic achievement were substantial starting from childhood. Education-specific noncognitive skills correlated strongly with achievement at the same age (*r* = 0.56 for the parent-rated factor and *r* = 0.84 for the teacher-rated latent measure), while the parent-rated and teacher-rated self-regulation factor correlated moderately with achievement (*r* = 0.37 and *r* = 0.25, respectively; Figure 1 and Supplementary Table 4).

#### Associations between individual noncognitive measures and general cognitive ability over development

Although noncognitive skills have traditionally been defined as all that is associated with educational attainment that is *not* cognitive skills^1^, constructs such as academic motivation, self-perceived ability, and interest have been found to be moderately linked to general cognitive ability^2^. We examined whether similar associations could be detected in our sample, where cognitive batteries had been administered at ages 7, 9, 12, and 16 (See Supplementary Note 1, and Supplementary Table 5 for the correlations between individual noncognitive traits and general cognitive ability).

Associations between *g* and individual self-reported education-specific noncognitive measures at the same age were mostly weak during childhood (ranging from r = 0.03, 95% CIs = 0.00, 0.05 for school enjoyment at age 9 to r = 0.26, 95% CIs = 0.26 for academic self-perceived ability at age 9). Associations with *g* increased to modest effect sizes during early adolescence (ranging between r = 0.21, 95% CIs = 0.19, 0.23 for academic interest at 12 and 0.37 for academic self-perceived ability at 12) and in some cases reached moderate effect sizes in late adolescence (e.g., r = 0.40, 95% CIs = 0.38, 0.43 for mathematics self-efficacy at age 16, and r = 0.36, 95% CIs = 0.34, 0.39 for academic self-concept at age 16). For other constructs associations remained weak across adolescence (e.g., r = 0.08, 95% CIs = 0.05, 0.11 for grit at age 16 and r = 0.11, 95% CIs = 0.08, 0.14 for curiosity; see Supplementary Table 5). Associations between *g* and parent and teacher-rated education-specific noncognitive measures were moderate even in childhood (e.g., r = 0.30, 95% CIs = 0.28, 0.33 for parent-rated academic interest at 9, r = 0.34, 95% CIs = 0.32, 0.37 for teacher-rated academic interest at 9, Supplementary Table 5). Associations between *g* and observed measures of low self-regulation remained stable over development and were characterized by weak effect sizes (from r = -0.19, 95% CIs = -0.21, -0.17 for parent-rated hyperactivity at 7 to r = -0.14, 95% CIs = -0.17, -0.11 for self-rated hyperactivity at 16; see Supplementary Table 5).

When considering cognitive and noncognitive factors, we found that general cognitive ability, constructed using factor analysis (see Supplementary Table 6 for factor loadings and model fit indices for all latent measures of general cognitive ability), was moderately associated with both education-specific noncognitive skills and self-regulation at all developmental stages. These associations were positive and ranged from .11 to .48 (Figure 1 and Supplementary Table 7). Similar to what we observed for academic achievement, associations between latent education-specific self-reported measures of noncognitive skills and general cognitive ability increased developmentally; effect sizes increased from weak (*r* = 0.11 at age 9) to moderate (*r* = 0.42 at age 16). On the other hand, associations between latent self-regulation self-report measures and general cognitive ability remained stable developmentally and showed modest effects (e.g., *r* = 0.19 at 9, and 0.20, at 16). Associations were stronger for parent- and teacher-reported measures of noncognitive skills, ranging between *r* = 0.24 for parent-rated self-regulation at age 7 and *r* = 0.48 for teacher-rated education-specific noncognitive skills at age 9.

### Supplementary Note 2

#### Genetic and individual-specific environments, but not family-wide environments, contribute to variation in noncognitive skills developmentally

##### The aetiology of individual noncognitive measures

At age 7 heritability estimates of parent and teacher-reported self-regulation ranged between 0.45 (95% CIs = 0.41,0.49) for parent-rated hyperactivity to 0.72 (95% CIs = 0.66, 0.74) for teacher rated conduct problems.

At age 9 the heritability (h^2^) of self-reported education-specific noncognitive measures were modest to moderate. Estimates ranged between h^2^= 0.13 (95% CIs = 0.00,0.26) for educational opportunities to h^2^ = 0.42 (95% CIs = 0.35,0.46) for academic self-perceived ability, and between h^2^ = 0.32 (95% CIs = 0.20, 0.43) and h^2^ = 0.44 (95% CIs = 0.33, 0.52) for self-reported measures of self-regulation, peer problems and conduct problems respectively. Heritability estimates were more substantial for parent and teacher-reported measures. Estimates for education related noncognitive measures ranged between h^2^ = 0.20 (95% CIs = 0.18, 0.22) and h^2^ = .82 (95% CIs = 0.75, 0.88) and between h^2^ = 0.43 (95% CIs = 0.31, 0.54) and h^2^ = 0.72 (95% CIs = 0.64, 0.76) for measures of self-regulation.

At age 12 heritability estimates for self-reported education-specific noncognitive measures ranged between h^2^ = 0.10 (95% CIs = 0.00, 0.22) for mathematics environment and h^2^ = 0.53 (95% CIs = 0.45, 0.57) for academic self-perceived ability. Estimates were moderate for self-reported self-regulation measures, ranging between h^2^ = 0.36 (95% CIs = 0.26, 0.42) for emotional problems and h^2^ = 0.46 (95% CIs = 0.36, 0.49) for hyperactivity. For teacher and parent-reported self-regulation measures, estimates were strong and ranged between h^2^ = 0.44 (95% CIs = 0.39, 0.47) for teacher-rated emotional problems to h^2^ = 0.74 (95% CIs = 0.71, 0.75) for parent-rated peer problems.

At age 16, education related noncognitive measures were moderately heritable, with estimates ranging between h^2^ = 0.20 (95% CIs = 0.05, 0.35) for time spent studying mathematics and h^2^ = 0.58 (95% CIs = 0.50, 0.61) for mathematics self-perceived ability. Self-reported measures of self-regulation were also moderately heritable, ranging between h^2^ = 0.35 (95% CIs = 0.31, 0.39) for conduct problems and h^2^ = 0.43 (95% CIs = 0.37, 0.46) for peer problems. See Supplementary Table 9 for all estimates.

##### The aetiology of latent noncognitive factors

The pie charts in Supplementary Figure 8 present the heritability, shared environmental, and nonshared environmental estimates for variation in latent factors of noncognitive skills, obtained running common pathway models. Genetic and nonshared environmental factors were found to be the main sources of variation in education-specific noncognitive skills at all ages, while the contribution of environmental factors shared between siblings raised in the same family was negligible. Heritability estimates for these latent dimensions of education-specific noncognitive skills ranged between h^2^ = 0.49 (95% CIs = 0.39, 0.60) for self-reported measures at age 9, and h^2^ = 0.87 (95% CIs = 0.83, 0.91) for parent-reported measures at age 9 (Figure 2 left panel and Supplementary Table 10 for the standardized estimates).

The aetiology of the latent dimensions of self-regulation differed from that of education-specific noncognitive dimensions, particularly when considering parent-reported self-regulation, which evinced substantial shared environmental effects at all ages, ranging between c^2^ = 0.15 (95% CIs =	0.06, 0.25) at age 7 and c^2^ = 0.29 (95% CIs =	0.18, 0.44) at age 9. On the other hand, when considering self-reported and teacher-reported measures, findings for self-regulation were in line with what was observed for education-specific measures, with heritability estimates ranging between h^2^ = 0.55 (95% CIs =	0.40, 0.72) for self-reported measures at age 9 and h^2^ = 0.75 (95% CIs =	0.69, 0.81) for teacher-reported measures at age 9, individual-specific environmental estimates ranging between e^2^ = 0.25 (95% CIs =	0.20, 0.31) for teacher-reported measures at age 9 and e^2^ = 0.36 (95% CIs =	0.32, 0.42) for self-reported measures at age 16, with negligible effects of family-wide environmental influences.

Supplementary Figure 8 also presents the genetic, shared, and nonshared environmental correlations between latent noncognitive dimensions and academic achievement at the same age. Genetic correlations (r_G_) ranged between moderate (r_G_ = 0.31) and strong (r_G_ = 0.96), with an average r_G_ of 0.53, while nonshared environmental correlations (r_E_) were substantially weaker, ranging between 0.03 and 0.62, with an average re of 0.24. Shared environmental correlations (r_C_) ranged from zero to unity but should be interpreted considering the general lack of shared environmental effects on self and teacher-reported noncognitive traits. When considering parent-reported measures of self-regulation, which evinced significant shared environmental variance, shared environmental correlations between noncognitive traits and achievement at the same age were 0.52 at age 7, 0.36 at age 9 and 0.27 at age 12 (Supplementary Tables 11 and 12).

We also considered the proportion of the phenotypic correlations between latent dimensions of noncognitive skills and academic achievement that was accounted for by genetic, shared, and nonshared environmental factors. Genetic factors accounted for between 65% (for parent-reported self-regulation at age 7) and 89% (for self-reported self-regulation at age 16) of those correlations. Shared environmental factors for between 0% (for self-reported self-regulation at age 9 and 16) and 27% (for parent-reported self-regulation at age 7); and nonshared environmental factors accounted for between 2% (for parent-reported self-regulation at age 12) and 22% (for self-reported education-specific noncognitive skills at age 9) of the associations.

**Supplementary Note 3: Sensitivity Analyses.**

a) ***Supplementary Analysis 1: Significance testing of the developmental increase in the PGS predictions of academic achievement.***

To test whether effect size estimates for the association between the Cog and NonCog PGS and academic achievement differed significantly over time, we fitted a series of structural equation models (Figure S3.1) in which we explicitly constrained to equality different sets of parameter estimates. This was done separately for the Cog and NonCog PGS, to allow for differences in their relative contributions jointly considered within the same model. Specifically, the models tested were as follows: Model a: all paramteres were freely estimated. Model b: all effects sizes for the Cog PGS (labelled ‘a’) and NonCog PGS (labelled ‘b’) were constrained to be equal over time (but different from each other). Model c: Only the NonCog PGS effects were constrained to equality while the Cog PGS were allowed to freely vary over development. Model d: opposite to model c, the Cog PGS effects were constrained to equality and NonCog PGS effects allowed to vary.

We then performed nested comparisons, with a chi-square difference test to select the best fitting model. As reported in Table S3.1, model d was favoured over the others, indicating that NonCog PGS effects differed over time while the Cog PGS effects could be constrained to equality over development.

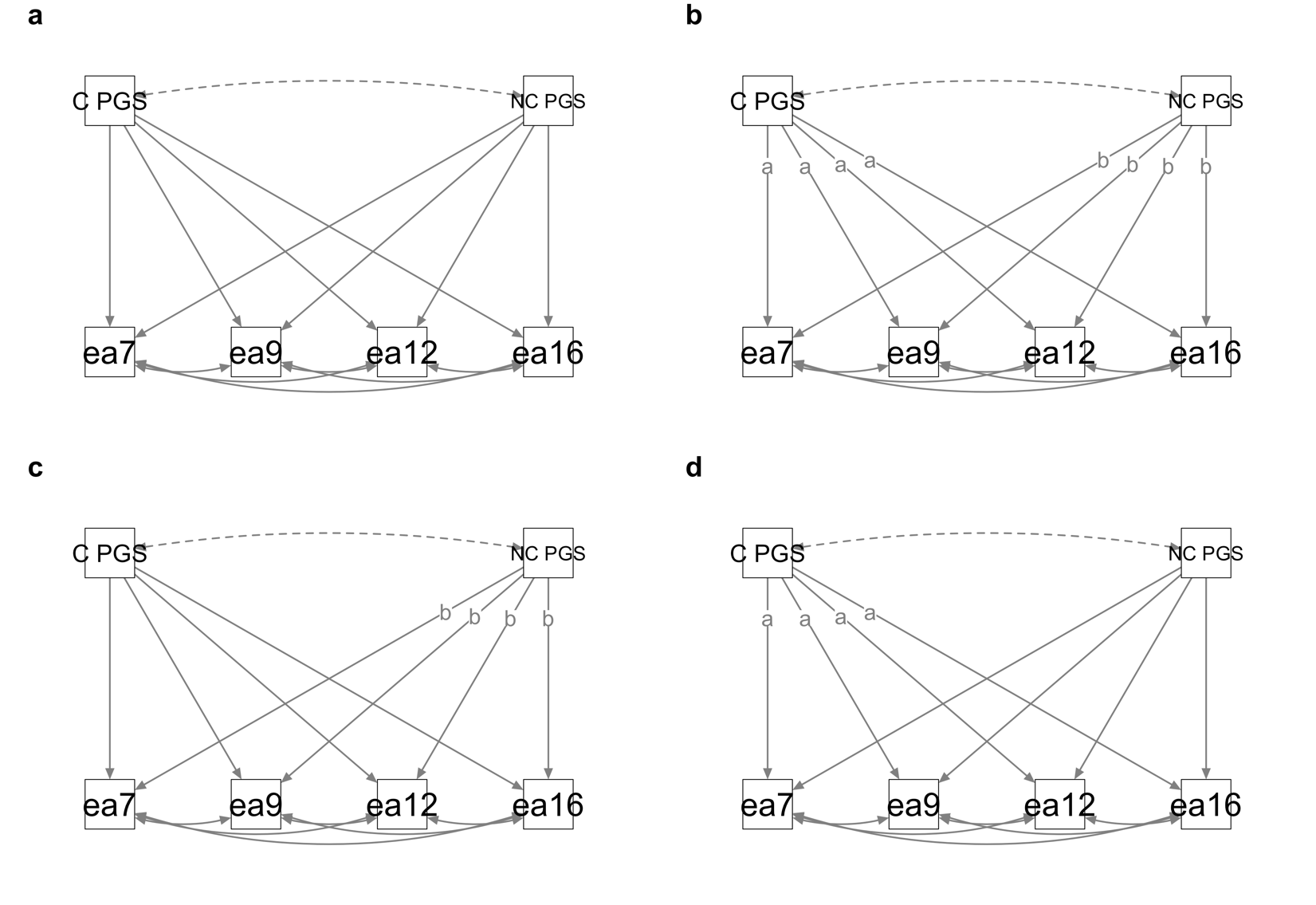

**Figure S3.1. Structural equation models testing differences between PGS effects over development.** C PGS = Cog polygenic score; NC PGS = NonCog polygenic score; ea = academic achievement

**Table S3.1. Model fit indices for the nested model comparisons between the four alternative models presented in Figure S.3.1.** AIC = Akaike information criterion; BIC = Bayesian information criterion; Chisq = chi square; Pr (>Chisq) = p value chi squared

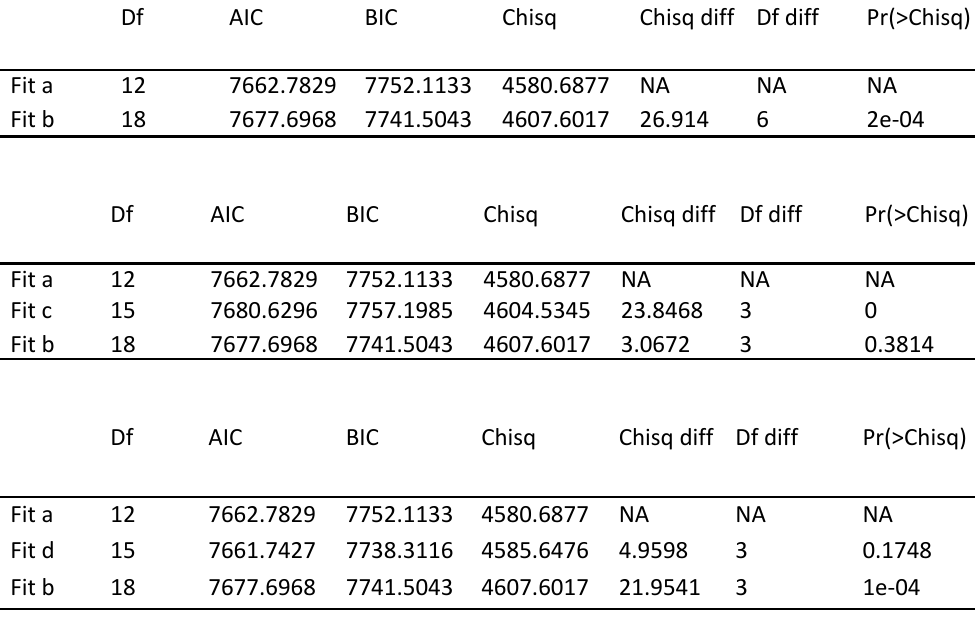

To further test whether this differences in effect size over time could be mainly attributable to a specific time point, such as for example age 16 being different if compared to other ages, we compared model d, with 3 further nested models where we allowed for an increasing number of time point to be constrained to equality (Figure S3.2). In model e, parameter estimates for the NonCog PGS prediction of achievement at ages 7 and 9 were constrained to equality. In model f the NonCog PGS predictions achievement at ages 7, 9 and 12 were constrained to equality. In model g the NonCog PGS predictions achievement at ages 7 and 12 were constrained to equality. As indicated by nested model comparisons reported in Table S3.2, we found that the PGS predictions at ages 9 and 16 could not be contained to equality without a decrease in model fit, indicating a significant difference.

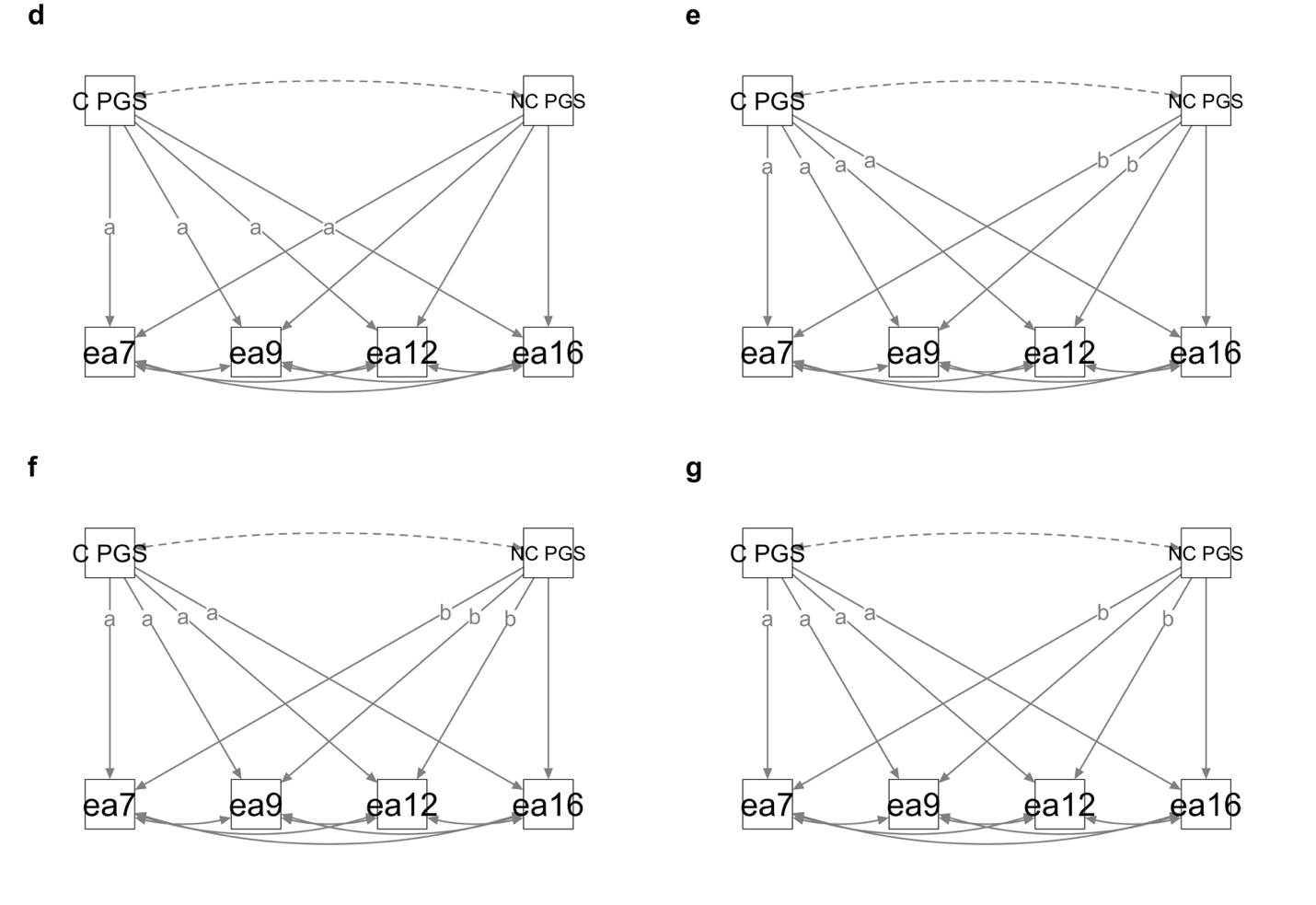

**Figure S3.2. Structural equation models testing differences between NonCog PGS effects over development.** C PGS = Cog polygenic score; NC PGS = NonCog polygenic score; ea = academic achievement

**Table S3.2. Model fit indices for the nested model comparisons between the four alternative models presented in Figure S.3.2.** AIC = Akaike information criterion; BIC = Bayesian information criterion; Chisq = chi square; Pr (>Chisq) = p value chi squared

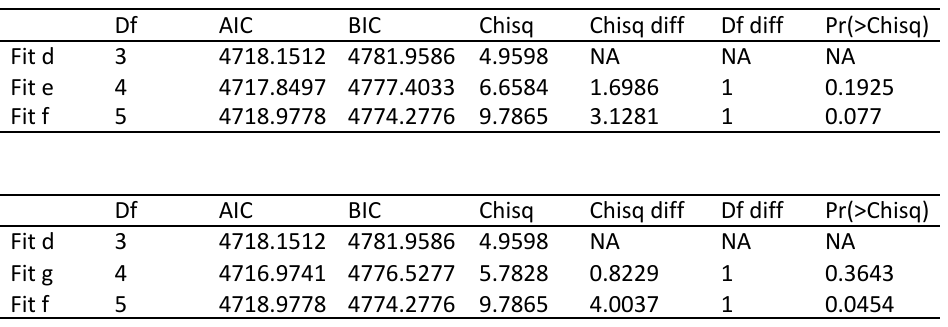

b) ***Supplementary Analysis 2: Cog and NonCog PGS predictions of academic achievement accounting for phenotypic g***

We also tested whether the observed increase in the NonCog PGS prediction of academic achievement was due to the possibility that the NonCog PGS simply captured more variance associated with cognitive ability rather than noncognitive skills. We addressed this potential issue by rerunning our Cog and NonCog PGS predictions of academic achievement including a measure of phenotypic general cognitive ability (g) in our multivariate models. For each PGS prediction we accounted for g measured at the same age as academic achievement. This very stringent test allowed us to examine whether the noncognitive PGS could still predict academic achievement at age 16 even after accounting for all the variance (not only genetic) shared with general cognitive ability. The results are presented in Table S3.1 below. Although, as expected, the effect of the prediction was attenuated, the NonCog PGS remained a significant predictor of variation in academic achievement at age 16.

**Table S3.3:** Noncognitive polygenic score prediction of academic achievement over development accounting for phenotypic general cognitive ability (g).

| **Measure of achievement** | **measure of g included** | **ß** | **Robust standard error** | **p value** |
| --- | --- | --- | --- | --- |
| Achievement age 7 | g age 7 | 0.06472661 | 0.01377481 | 2.62E-06 |
| Achievement age 9 | g age 9 | 0.08331642 | 0.01697248 | 9.16E-07 |
| Achievement age 12 | g age 12 | 0.07146178 | 0.01742392 | 4.11E-05 |
| Achievement age 16 | g age 16 | 0.1629107 | 0.01516482 | 6.42E-27 |

Note: All multivariate regression included the following predictors (in addition to the NonCog PGS and g): Cog PGS, the first 10 principal components of ancestry, genotyping chip. All variables were residualized for age and sex and the residuals were standardized before regression analyses.

c) ***Supplementary Analysis 3: Significance testing of the developmental increase in the PGS predictions of academic achievement adjusting for SES.***

We conducted the same analyses reported in Supplementary Note 3, this time adjusting for socio-economic status. Adjusting the predictors and outcomes for SES did not change our conclusion: while Cog PGS effects did not significant change over time, NonCog PGS did. Model fit indices are reported in Table S3.4

**Table S3.4. Model fit indices for the nested model comparisons between the four alternative models presented in Figure S.3.2, adjusting for phenotypic socioeconomic status .** AIC = Akaike information criterion; BIC = Bayesian information criterion; Chisq = chi square; Pr (>Chisq) = p value chi squared

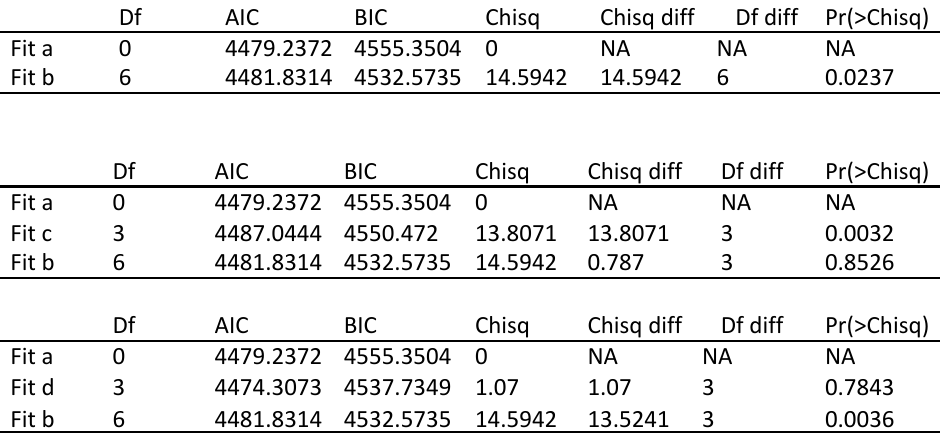

**Figure 1.** **Contemporaneous correlations between observed noncognitive measures and academic achievement from age 7 to 16.** **Panel a.** presents the correlations between academic achievement and education related noncognitive measures. **Panel b.** shows the correlations between academic achievement and observed measures of self-regulation. The length of each bar indicates the size of the correlation and the error bar 95% confidence intervals, the numbers on the right indicate the different ages.

**Figure 1a.** Education related noncognitive measures. spa = self-perceived ability; satisfaction = classroom satisfaction; opportunity = educational opportunity; interest = academic interest; enjoyment = enjoyment of learning; lit environment = literacy environment; attitudes = attitudes towards learning.

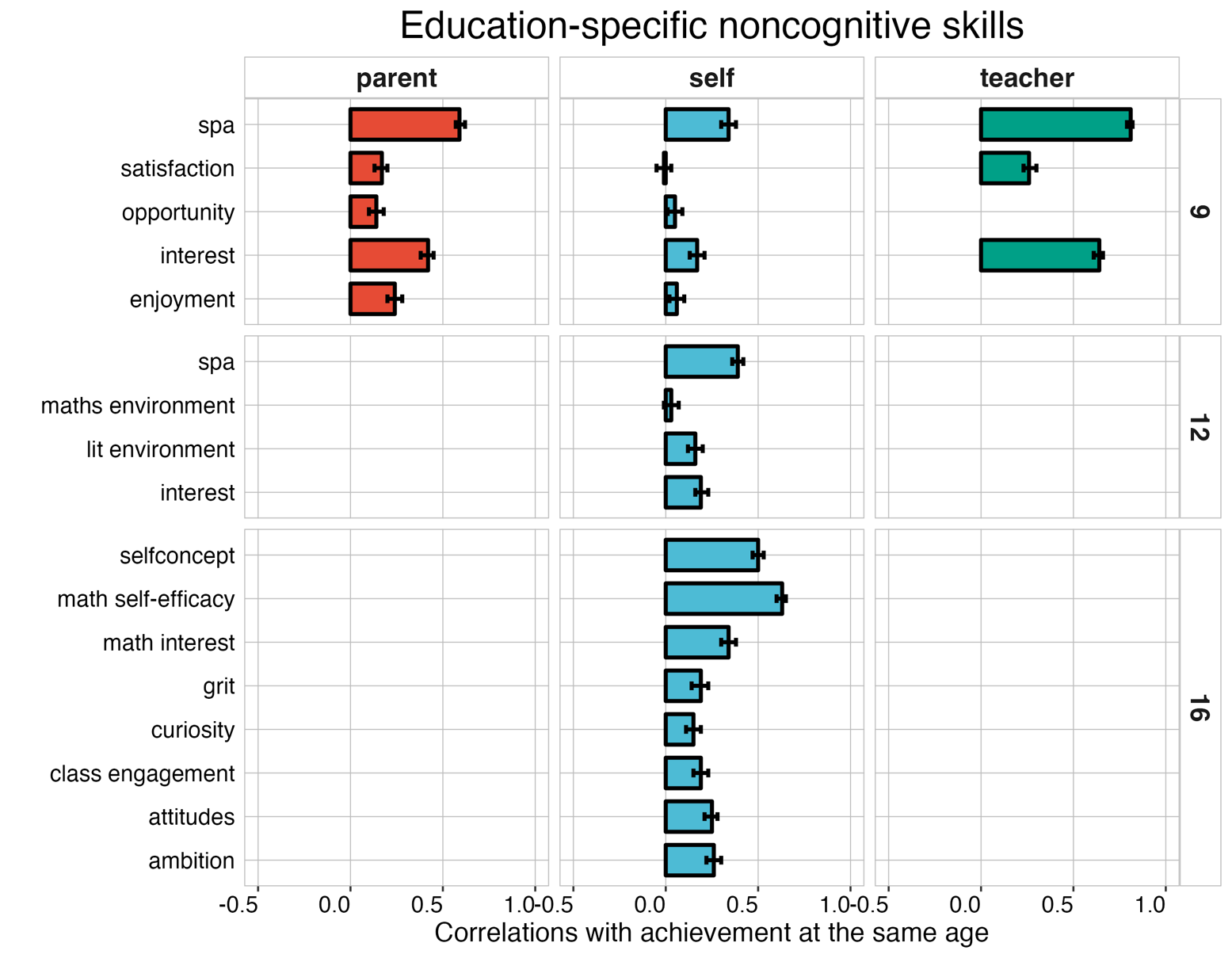

**Figure 1b. Self-regulation measures.** Prosocial = prosocial behaviour; peerproblems = peer problems; conduct = conduct problems.

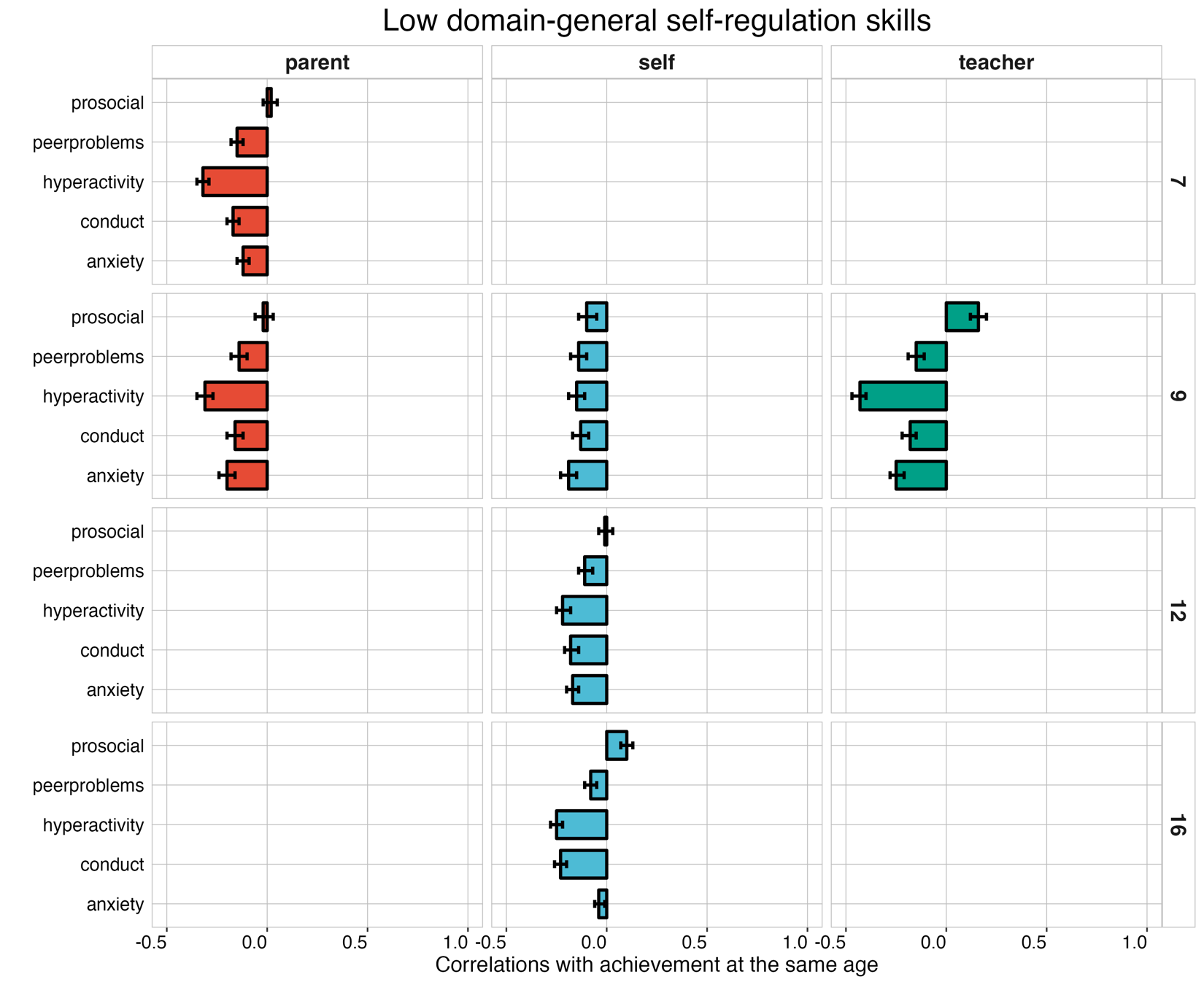

**Figure 2.** Associations between latent noncognitive measures, academic achievement, and general cognitive ability. The top panel shows the associations between latent education-specific noncognitive measures and academic achievement at the same age (red bars), cognitive ability at the same age (blue bars), and academic achievement after accounting for general cognitive ability at the same age using a multiple regression framework (yellow bars). Each bar indicates the effect size of standardized regression coefficients and the error bars the 95% confidence intervals around each estimate. The bottom panel shows the same associations for latent measures of self-regulation over development. The different panels show developmental trends for different reporters, from left to right, parent-reported, teacher-reported, and self-reported noncognitive dimensions.

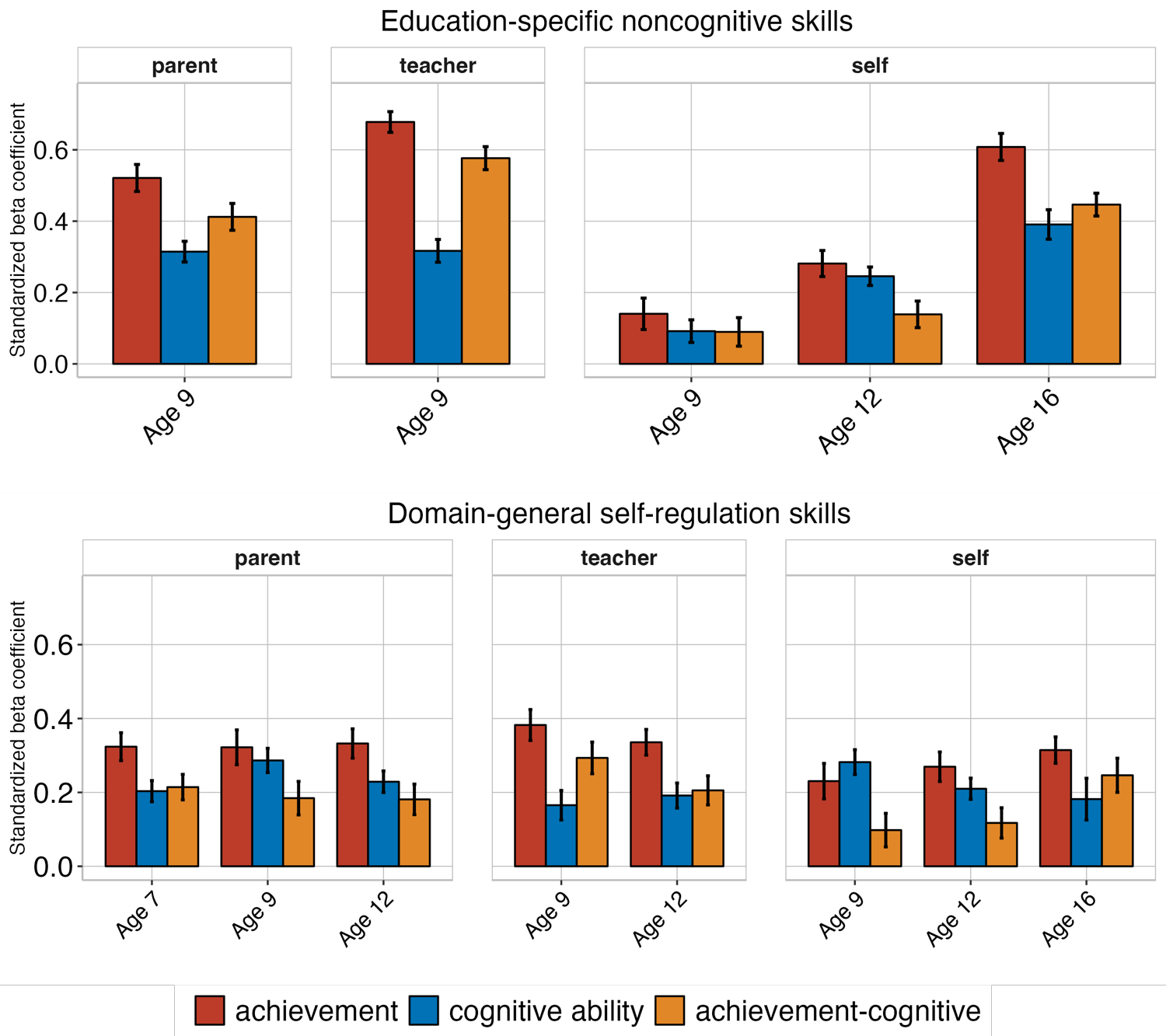

**Figure 3. Heritability (a^2^), shared environmental (c^2^), and nonshared environmental (e^2^) estimates for all observed measures of education-specific noncognitive skills from age 9 to 16.** **Panel A** shows the estimates for self-rated measures which were collected when the children were 9, 12, and 16. **Panel B** presents the estimates for parent-rated education-specific noncognitive measures, which were collected when the twins were 9 years old. **Panel C** presents the estimates for teacher-rated education-specific noncognitive measures, collected when the twins were 9 years old. Error bars are 95% confidence intervals. The numbers at the top indicate the different ages.

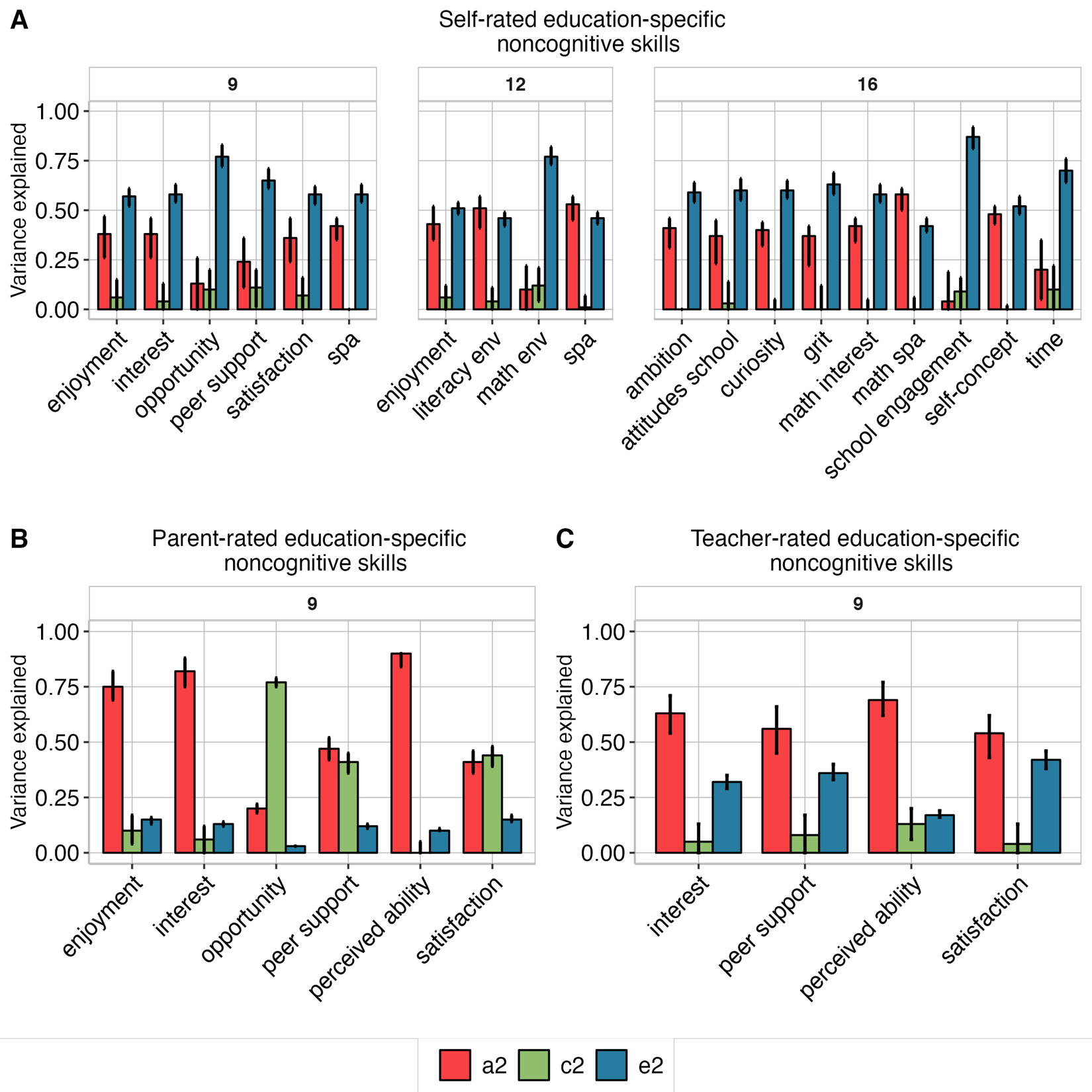

**Figure 4. Heritability (a^2^), shared environmental (c^2^), and nonshared environmental (e^2^) estimates for all observed measures of domain-general self-regulation.** Measures were collected at ages 7, 9, 12 and 16 and ratings were obtained from parents (top row), teachers (bottom row) and self-rated by the twins (middle row). Error bars are 95% confidence intervals. The numbers at the top indicate the different ages and the panels on the right the different raters (parents, teachers, and self).

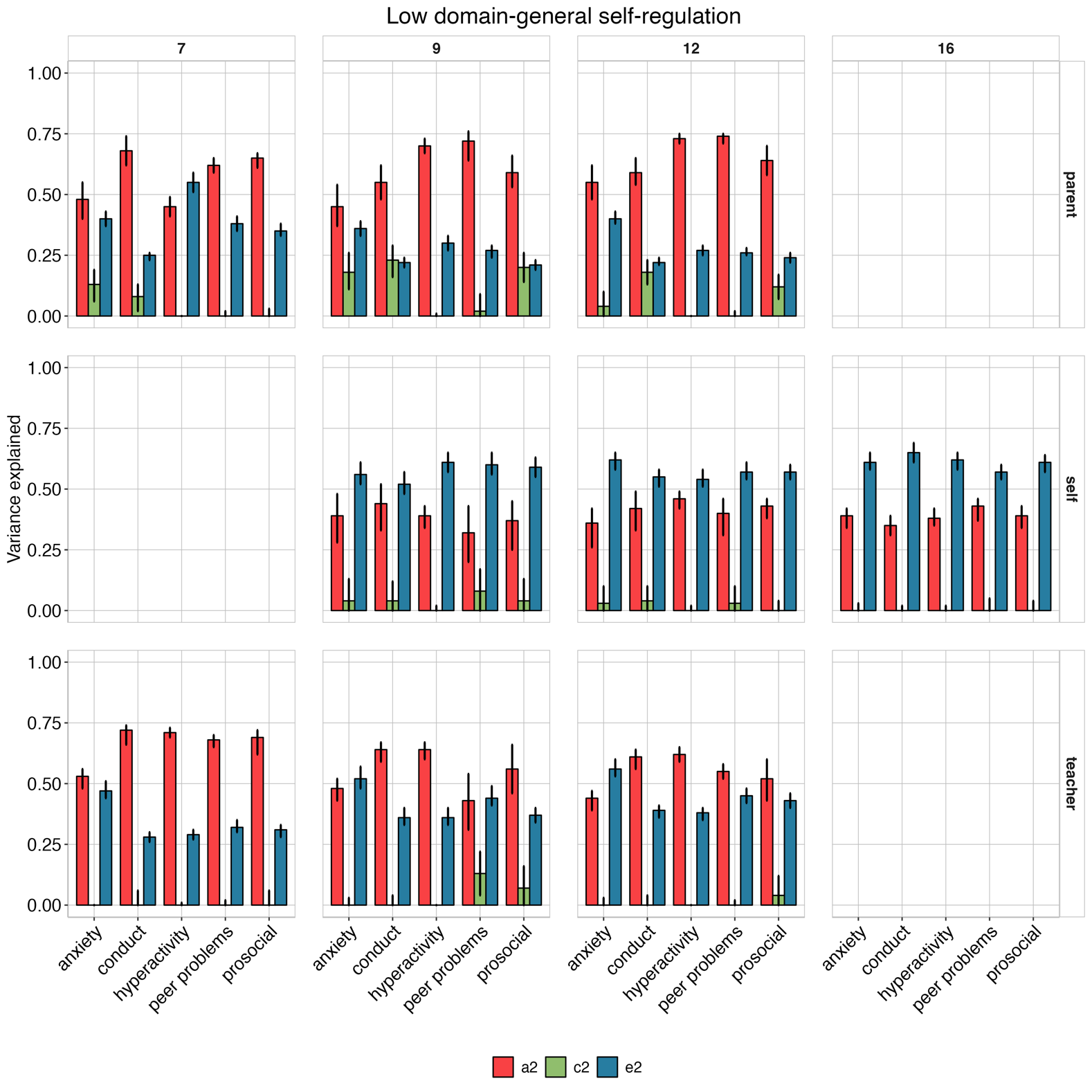

**Figure 5. Heritability (a^2^), shared environmental (c^2^), and nonshared environmental (e^2^) estimates for all observed measures of academic achievement from age 7 to 16.** Error bars are 95% confidence intervals. The numbers at the top indicate the different ages.

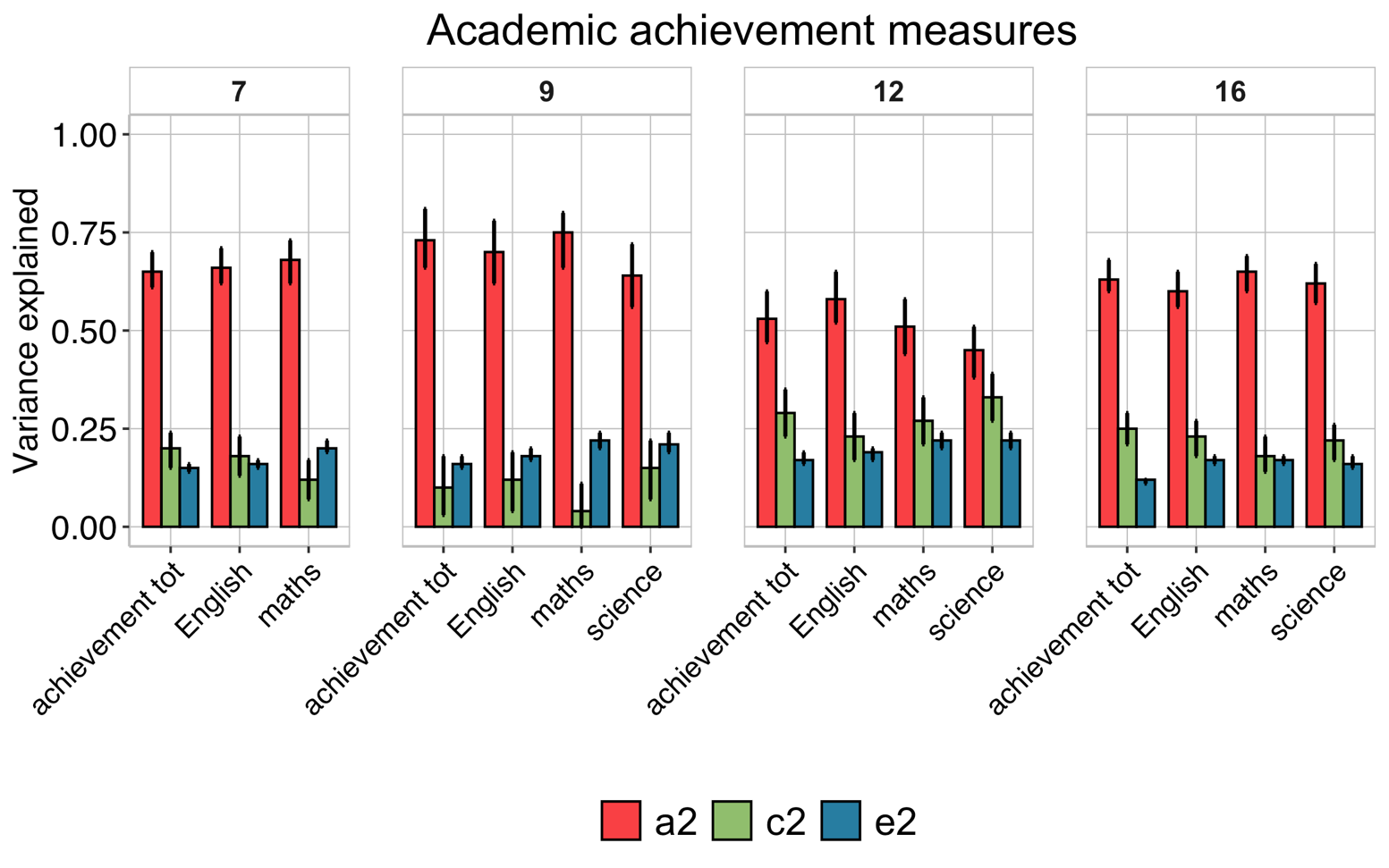

**Figure 6. Heritability (a^2^), shared environmental (c^2^), and nonshared environmental (e^2^) estimates for all observed measures of cognitive ability from age 7 to 16.** Error bars are 95% confidence intervals. The numbers at the top indicate the different ages. g = total general cognitive ability composite.

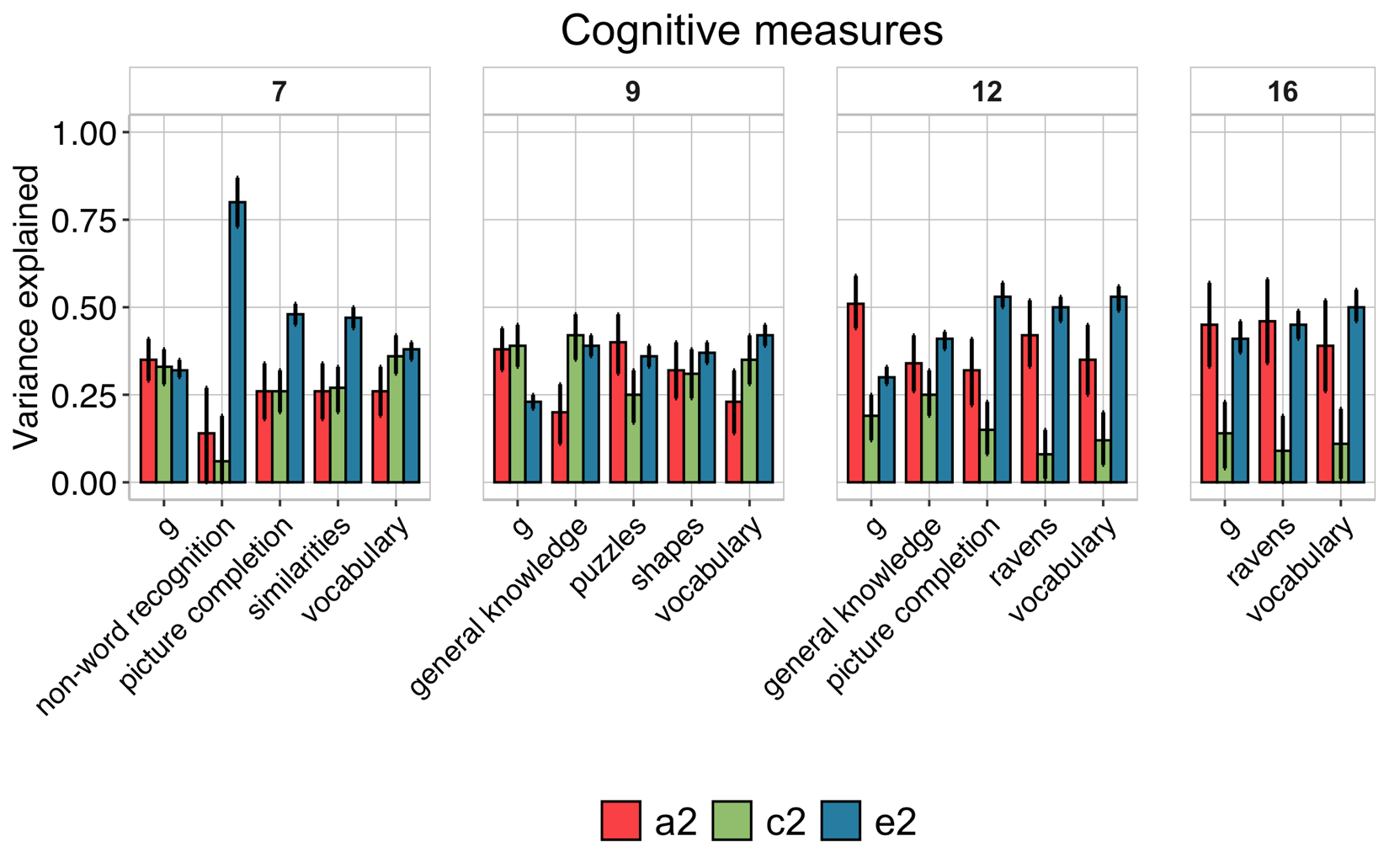

**Figure 7. Heritability (a^2^), shared environmental (c^2^), and nonshared environmental (e^2^) estimates for all observed measures of early academic abilities at age 4.** Error bars are 95% confidence intervals.

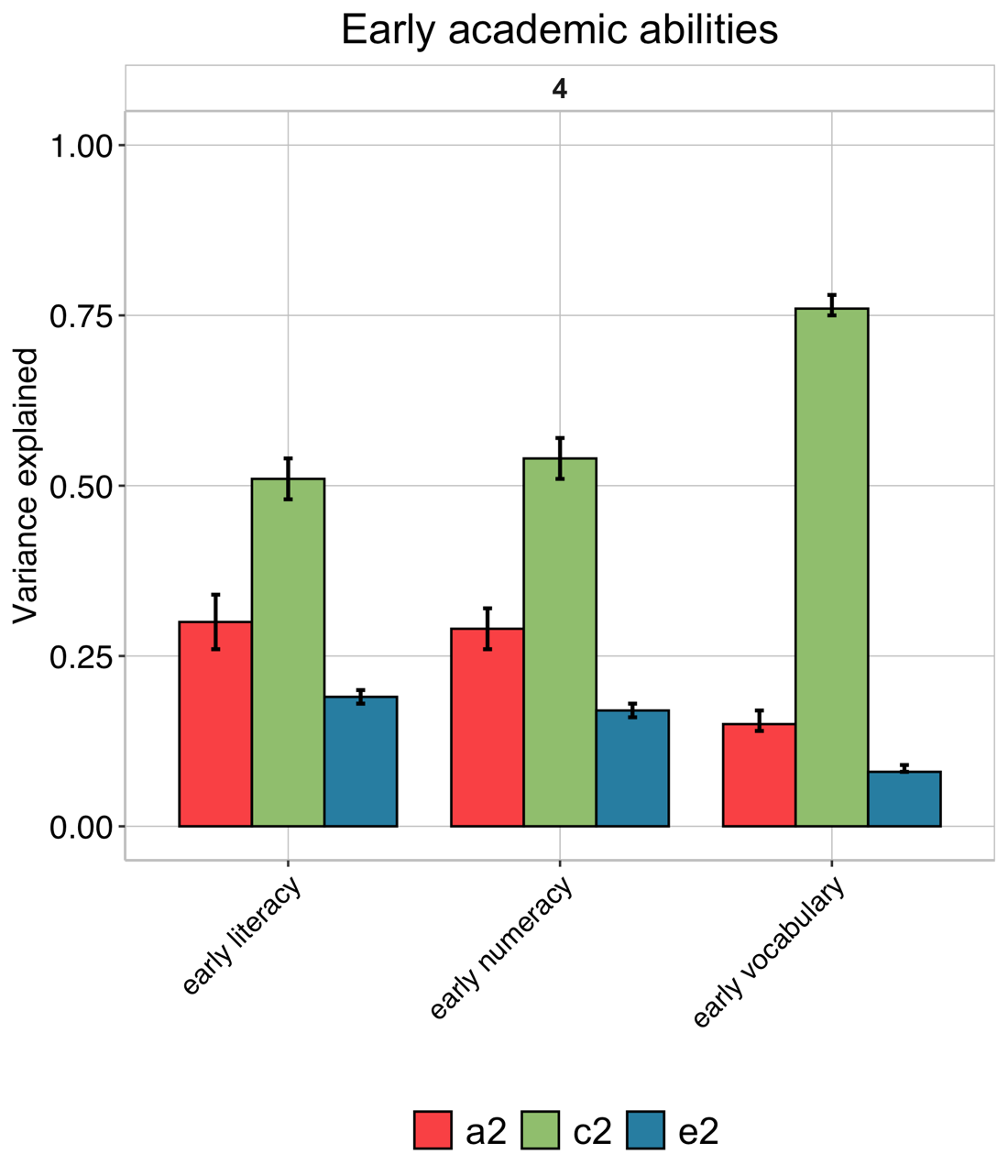

**Figure 8.** **Genetic and environmental influences on latent noncognitive dimensions and on their associations with academic achievement at the same age.** The pie charts represent the proportions of variation in the latent dimensions of education-specific noncognitive skills (left) and self-regulation (middle and right) accounted for by genetic (A) shared environmental factors (C) and nonshared environmental factors (E). The balloon plots next to each pie chart depict the size of the genetic (in red), shared environmental (in green), and nonshared environmental (in blue) correlation between each latent noncognitive dimension and academic achievement at the same age. P = parent-reported, T = teacher-reported, S = self-reported.

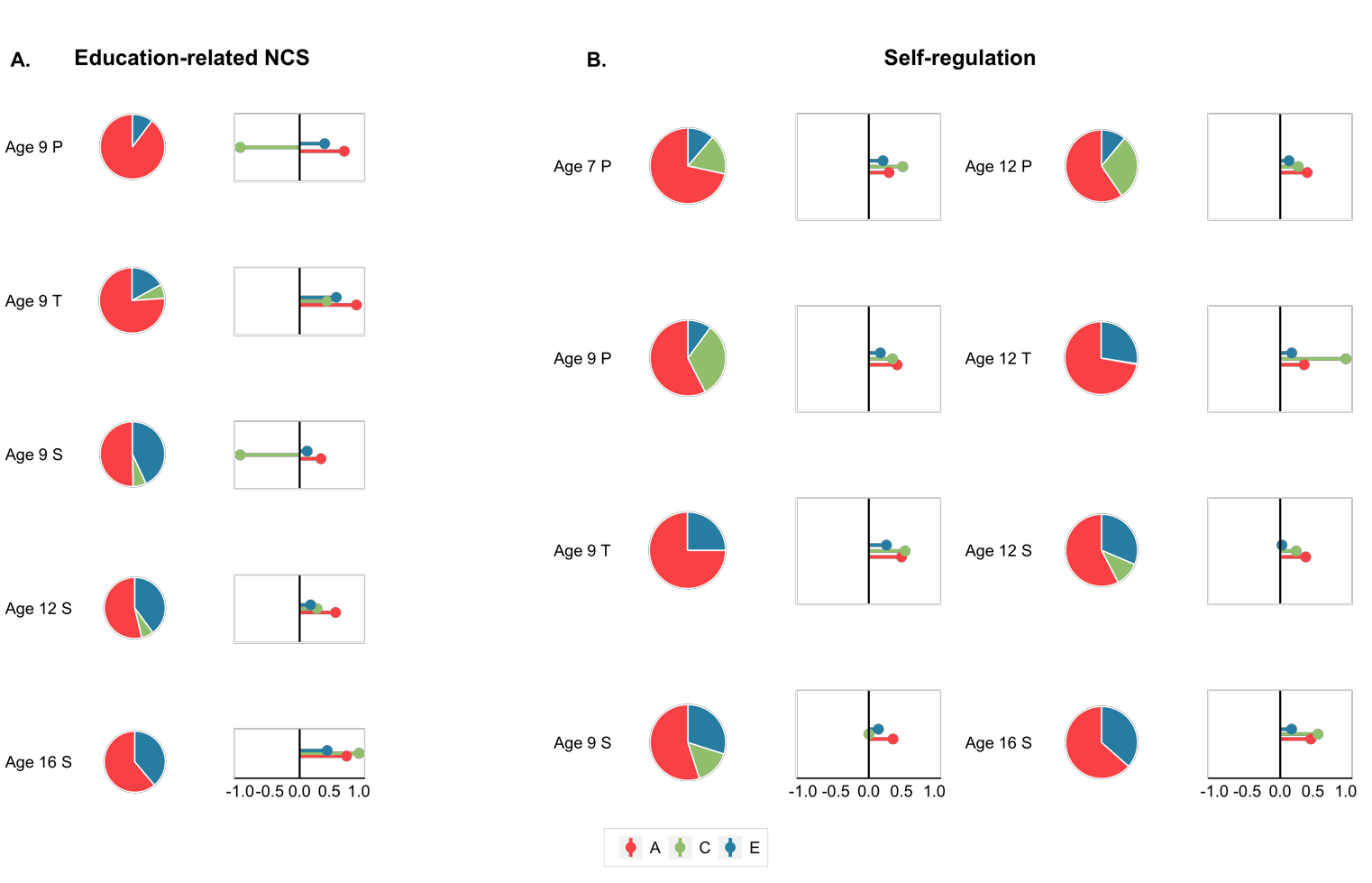

**Figure 9. Contribution of genetic and environmental influences on cognitive and noncognitive skills to academic achievement at the same age.** Each bar presents the outcomes of a trivariate Cholesky decomposition. The proportion of variance in academic achievement is partitioned into genetic (orange), shared environmental (green), and nonshared environmental (blue) influences. The lighter shadings indicate the proportion of variance in academic achievement accounted for by genetic (A cog) and environmental (C cog and E cog) variance in cognitive skills. The darker shadings indicate the proportion of variance in academic achievement accounted for by genetic (A noncog−cog) and environmental (C noncog−cog and E noncog−cog) variance in noncognitive skills independent of genetic and environmental effects on cognitive skills. The darkest shadings indicate genetic and environmental effects on academic achievement independent of genetics and environmental variance in cognitive and noncognitive skills. 95% Confidence intervals for all estimates are presented in Supplementary Tables 12 and 13.

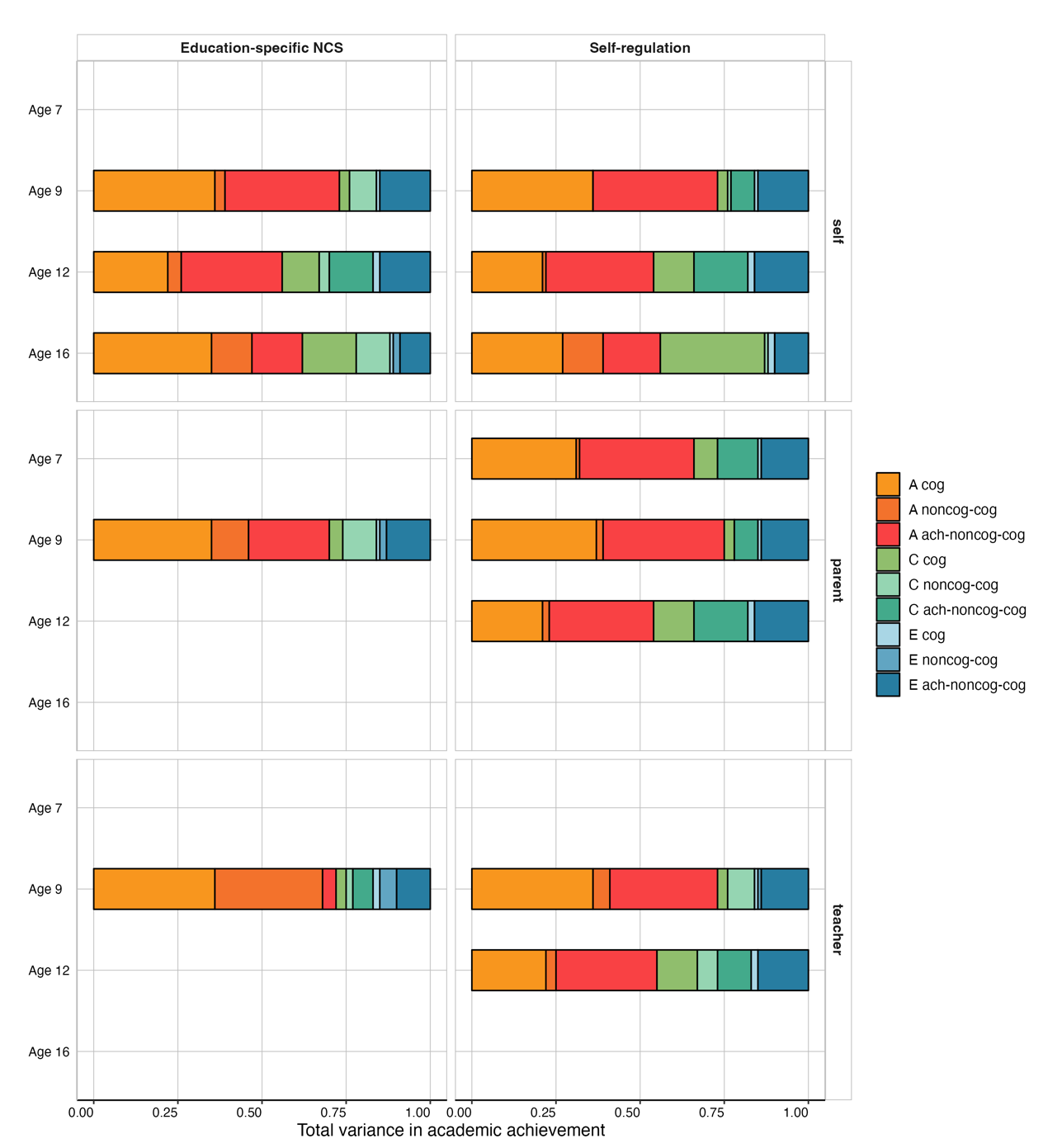

**Figure 10. Genetic correlations.** Correlations (Rgs) are presented for the new cognitive (Cchol) and noncognitive (NCchol) factors obtained with the new extension of the GWAS-by-subtraction model. These are compared against the correlations obtained for the original Cog (C) and Noncog (NC) factors obtained using the GWAS-by-subtraction (Damange et al. 2021). SES = socioeconomic status.

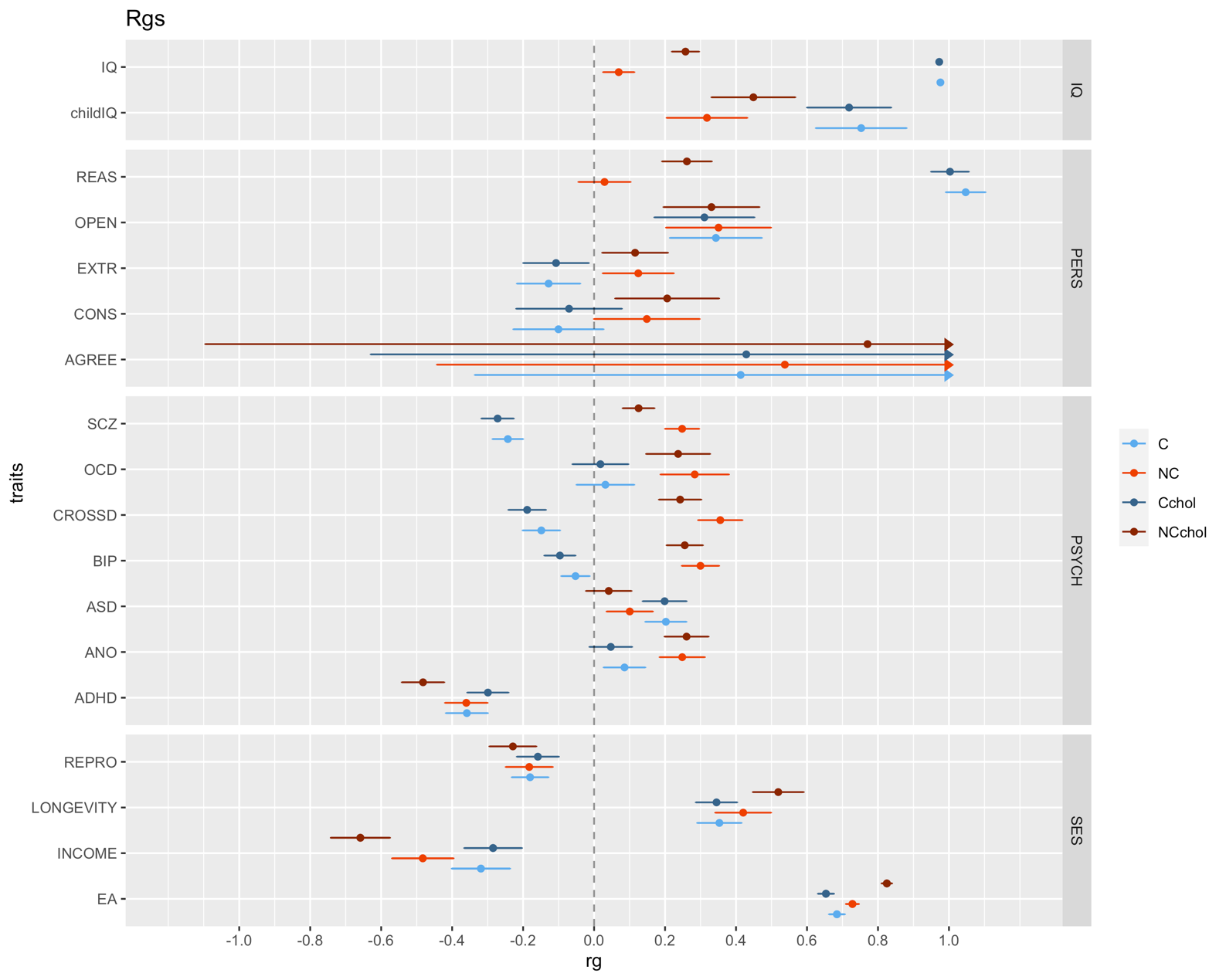

**Figure 11. Cognitive and noncognitive polygenic score (PGS) predictions of noncognitive, academic achievement and family socio-economic status phenotypes over development.** Panel a. shows the PGS prediction of education-specific and self-regulation latent phenotypes from age 7 to 16. Panel b. shows the PGS prediction of early indicators of academic achievement at age 4. Panel c shows the PGS prediction of family socioeconomic status measured at first contact (ages 0-2) and again when the twins were 16 years old. The light blue and orange dots represent the PGS calculated based on the summary statistics for the original GWAS-by subtraction model (Demange et al., 2021), and the dark blue and red dots represent the PGS calculated from the summary statistics obtained with our extension of the model (Cog and NonCog; **Supplementary Table 14** and **Figure 3A**).

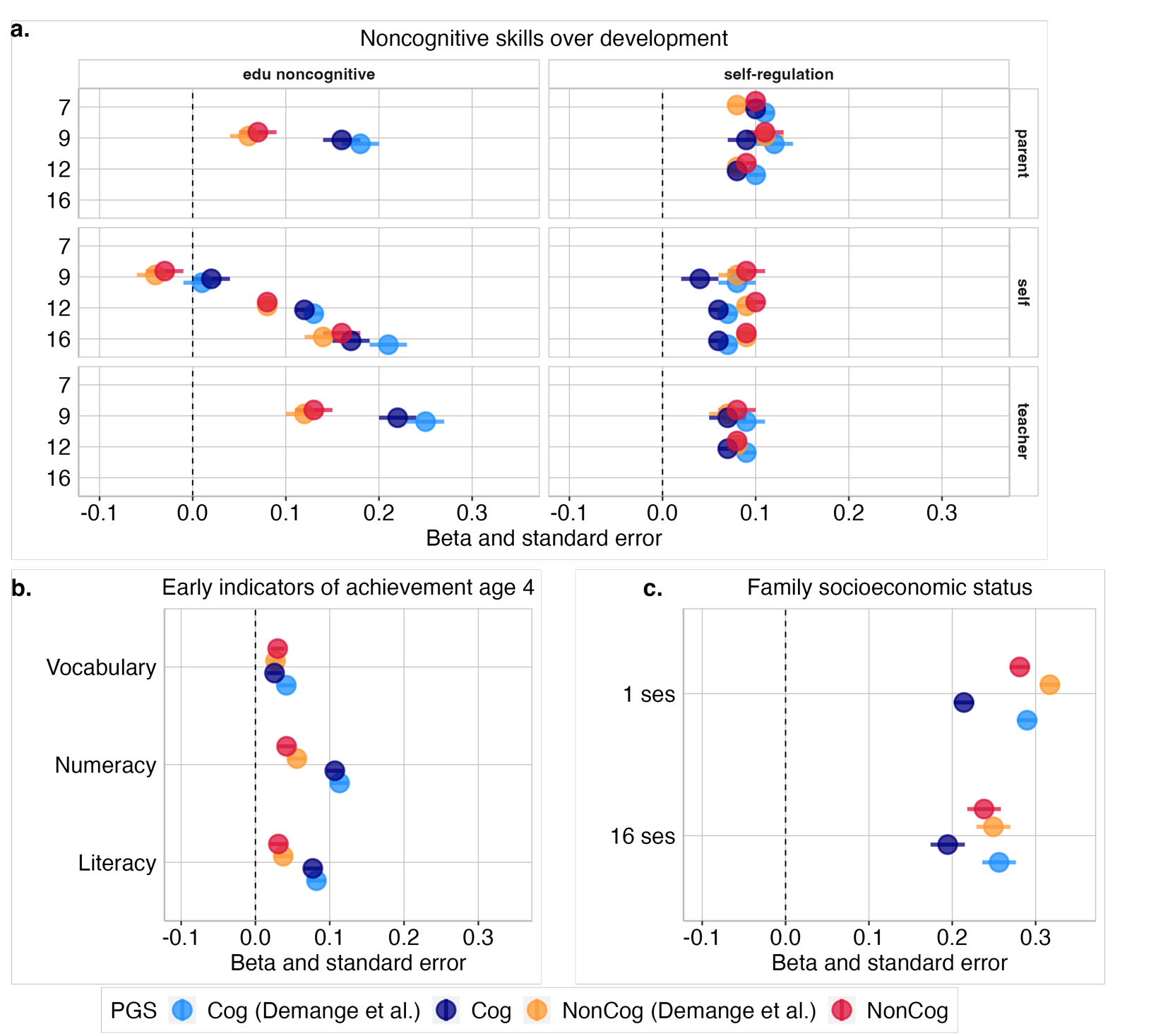

**Figure 12. Cognitive and Noncognitive polygenic score (PGS) predictions of cognitive phenotypes over development.** PGSs were derived from the original (Demange et al., 2021) and the extended GWAS-by-subtraction model (Cog and NonCog).

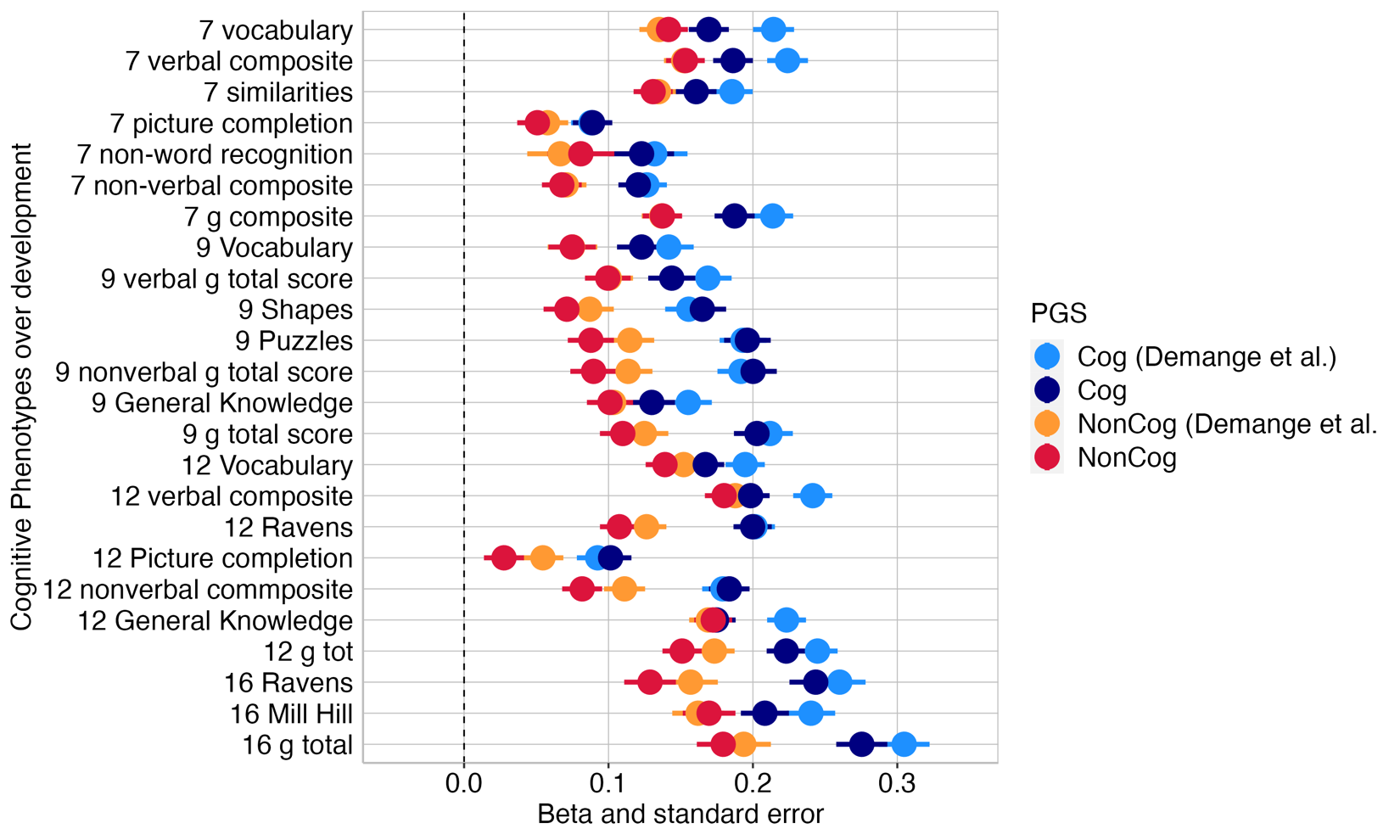
